# Encoding activity predicts distinct recognition signals during working and long-term memory in the human hippocampus

**DOI:** 10.64898/2026.10.21.759719

**Authors:** Catherine W. Tallman, John T. Wixted, Peter N. Steinmetz

## Abstract

Neurocomputational models predict that episodic memories are represented by sparse, pattern-separated hippocampal codes. Recognition within working memory capacity may instead rely on overlapping cortical codes. We analyzed human intracranial microwire recordings during a continuous recognition task to test whether retrieval demand and spiking activity at encoding determine which neural signal is expressed. We show that relative increases in firing from pre-stimulus to post-stimulus onset at encoding predicted an item-specific signal that was present only in the hippocampus and only during long-term memory retrieval of remembered items. In contrast, consistently elevated firing predicted a generic signal that distinguished remembered from novel items, was present in the hippocampus during both working and long-term memory retrieval, and extended to the anterior cingulate and prefrontal cortices during working memory. These findings are consistent with a new theory that distinguishes working memory from long-term memory in terms of the neural coding scheme accessed at retrieval.

## Introduction

Human recognition memory can be supported by different forms of neural representation. The classic distinction between short-term and long-term memory was motivated in large part by neuropsychological evidence showing that medial temporal lobe damage profoundly impairs long-term declarative memory while leaving limited short-term maintenance relatively preserved ^1–4^. For example, Patient H.M.’s recognition memory for digits and pure tones over retention intervals of up to approximately 5 seconds was well within the normal range ^5^, yet hippocampal involvement is not determined by retention interval alone. Hippocampal contributions have also been observed in tasks conventionally classified as working memory, including under high memory loads and in some tasks requiring relational or high-resolution information ^6–9^. What determines which neural coding system supports recognition under different memory demands?

A recently updated Atkinson-Shiffrin framework linking short- and long-term memory to Complementary Learning Systems (CLS) theory ^10,11^ offers a specific account ^12^. Information that remains directly accessible over the short term is proposed to be supported by a transient, task-ready cortical state instantiated by overlapping neural codes, such that shared features of different items are represented by overlapping neuronal populations. This cortical state is vulnerable to interference and is distinct from the slowly consolidated cortical representations emphasized in standard CLS. In parallel, the hippocampus can form a sparse, pattern-separated index of the recent episode. While the cortical state remains sufficiently accessible, performance need not depend on retrieval through that index. As subsequent input reconfigures and blurs the cortical state ^13^, however, hippocampal retrieval should become increasingly consequential for recovering the earlier episode.

Cortical coding therefore contributes under both working- and long-term-memory conditions. During working memory, the recently configured cortical state may support performance directly while during long-term-memory retrieval, a hippocampal index can recover the episode by reinstating the associated cortical representation. Thus, hippocampal indexing provides an additional route for recovering the relevant cortical state rather than replacing cortical representation. These coding schemes are also implemented in CLS: dentate gyrus-CA3 circuitry is proposed to form sparse, pattern-separated representations that preserve the discriminability of similar experiences despite interference ^10,11,14–18^, whereas neocortical coding is more distributed and overlapping, allowing shared features and structure to be represented across experiences ^10,11^. Overlapping cortical codes are well suited to integrating and comparing currently accessible information, whereas sparse hippocampal codes are well suited to individuating specific episodes.

Intracranial recordings from epilepsy patients completing recognition memory tasks revealed two neural signals that provide a means of testing this account. A generic recognition signal, expressed as greater mean firing for previously encountered than novel items, has been observed across the hippocampus, amygdala, anterior cingulate cortex (ACC), and prefrontal cortex (PFC) ^19–23^. Thus, it is broadly distributed and compatible with cortical representations contributing whether directly accessible or reinstated through hippocampal retrieval. A second, item-specific signal has a markedly different profile: hippocampal spike-count distributions are more positively skewed for studied items than for novel items ^21^, reflecting a small number of disproportionately high-firing neuron-by-item responses. This hippocampal-selective signature has properties expected of sparse, item-specific episodic coding. One key question is whether it is selective to retrieval of individual episodic memories while the generic signal remains evident across levels of retrieval demand.

The two signals may also be associated with different neural states at encoding. In an independent delayed study-test dataset, the item-specific hippocampal signal was selectively observed for subsequently remembered items ^24^ associated with a transition from relatively low pre-encoding firing to relatively high firing during encoding, broadly consistent with neuronal-allocation models ^25^. Consistently high firing before and during encoding, by contrast, resembles the broader pattern of elevated encoding-related activity associated with subsequent memory across ERP, fMRI, and intracranial recording studies ^26–29^. We therefore asked whether these distinct encoding states differentially predicted expression of the item-specific and generic signals.

Here, we retrospectively analyzed intracranial microwire recordings previously used to identify these signals, examining encoding state and retrieval demand within the same participants, task, and neuronal populations. Patients with epilepsy performed a continuous-recognition task in which words were repeated after 0, 1, 3, 7, 15, or 31 intervening items. Short lags fell within typical working memory capacity, whereas longer lags introduced substantially more intervening information and thus a greater likelihood that the task-ready cortical state would be reconfigured and blurred. We therefore treated short- and long-lag conditions as imposing working memory and long-term-memory retrieval demands, respectively.

We predicted that the item-specific signal would be associated with a transition from relatively low pre-encoding firing to relatively high firing during encoding and tested whether the generic signal was likewise related to encoding state. Most critically, if the item-specific signal reflects retrieval of a sparse, pattern-separated episodic representation, it should become more pronounced under long-term-memory than working memory retrieval demands, whereas the generic recognition signal should remain evident across both conditions.

## Results

Patients with medically intractable epilepsy undergoing clinical monitoring completed a continuous recognition memory task in which single words were presented twice, with a variable number of intervening items. We analyzed 55 recording sessions from 34 patients implanted with depth electrodes targeting the hippocampus, amygdala, anterior cingulate cortex (ACC), and prefrontal cortex (PFC). Spike counts were measured 200-1000 ms after word onset and normalized relative to each neuron’s session-wide baseline activity, measured -1000 to -200 ms relative to stimulus onset.

Retrieval demand was defined by the number of intervening items between the first and second presentation of each word. Words with short lags (0, 1, and 3 intervening items) were categorized as working memory (WM), whereas words with longer lags (7, 15, and 31 intervening items) were categorized as long-term memory (LTM). Longer-lag conditions exceeded typical working memory capacity limits of approximately four items ^30,31^ and were predicted to require episodic retrieval beyond active maintenance. Each repeated item was also categorized by the relative level of spiking activity elicited directly before and during stimulus presentation at encoding: increased (Low-High), consistently elevated (High-High), decreased (High-Low), and consistently low (Low-Low) firing.

To distinguish generic recognition from episodic item-specific retrieval, we compared differences in the first four moments of the repeated- and novel-item spike count distributions (mean, standard deviation, skewness, and kurtosis). Each moment was calculated from the pooled neuron-by-item distributions across sessions, where each observation represented the normalized spike count response of one neuron to one item presentation, rather than subject- or neuron-level summary distributions. Repeated-item distributions were constructed separately within each brain region (hippocampus, amygdala, ACC, and PFC) and for each item, categorized by retrieval-demand condition (working memory and long-term memory), encoding-defined firing condition (Low-High, High-High, High-Low, and Low-Low), and subsequent-memory outcome (remembered and forgotten). Novel-item distributions within each region served as the common comparison distribution across the repeated-item conditions. Statistical significance was evaluated using bootstrap tests with Benjamini-Hochberg correction for multiple comparisons (see Methods).

Primary analyses focused on remembered Low-High and High-High repeated items. Forgotten-item trials were substantially less frequent, limiting statistical power for reliable comparisons. Corresponding analyses of forgotten items are reported in Tables 1-2 and Tables S2-S5, and analyses of the High-Low and Low-Low conditions are reported in Supplemental Material (Tables S2-S5; Figs. S1-S2). Total trial counts and neuron-by-item sampling are reported in Table S1. Parallel analyses were conducted in an independent human single-neuron dataset collected during delayed study-test of objects, scenes, and animals ^32,33^. These data were acquired in a different laboratory and patient sample using different spike-sorting procedures and visual rather than verbal memoranda. The item-specific analysis presented in Figure 1A was reported previously ^24^, but the independent dataset here was used to test whether the item-specific memory signal generalized across task design, stimulus modality, and signal-processing approach.

**Table 1.** Item-specific memory signal (skewness) across relative levels of spiking at encoding and brain regions in the continuous recognition task for remembered and forgotten items. Repeated and novel distributions consisted of normalized single-neuron spike counts, with each observation representing the response of one neuron to one item. Repeated items were categorized according to the relative levels of spiking immediately before and during encoding. Primary analyses are shown for relative increases in firing (Low-High) and consistently elevated firing (High-High); results for all four firing patterns are reported in Supplementary Tables S2-S5. Retrieval demand was manipulated by lag, with long-term memory (LTM) corresponding to lags 7, 15, and 31 and working memory (WM) corresponding to lags 0, 1, and 3. ΔH-Region reflects the difference between the hippocampal repeated-novel effect and the corresponding repeated-novel effect in the comparison region. Statistical comparisons were assessed using two-sided bootstrap tests (10,000 resamples). Bolded *P* values survived Benjamini-Hochberg false-discovery-rate correction.

| Skewness |  |  |  |  |  |  |  |  |  |  |  |  |  |  |  |  |  |  |  |  |
| --- | --- | --- | --- | --- | --- | --- | --- | --- | --- | --- | --- | --- | --- | --- | --- | --- | --- | --- | --- | --- |
|  |  |  | Hippocampus |  |  | Amygdala |  |  |  |  | Anterior Cingulate Cortex |  |  |  |  | Prefrontal Cortex |  |  |  |  |
| | | Firing Pattern | Repeated | Novel | *P* | $\Delta H-A$ | *P* | Repeated | Novel | *P* | $\Delta H-AC$ | *P* | Repeated | Novel | *P* | $\Delta H-PF$ | *P* | Repeated | Novel | *P* |
| Remembered | LTM | Low-High | 4.17 | 2.08 | **0.006** | 2.26 | **0.004** | 1.85 | 2.02 | 0.263 | 1.58 | **0.040** | 2.97 | 2.46 | 0.123 | 2.15 | **0.016** | 2.59 | 2.65 | 0.907 |
|  |  | High-High | 1.79 | 2.08 | 0.505 | -0.09 | 0.854 | 1.82 | 2.02 | 0.138 | -0.57 | 0.284 | 2.74 | 2.46 | 0.377 | 0.02 | 0.955 | 2.34 | 2.65 | 0.428 |
|  | WM | Low-High | 2.17 | 2.08 | 0.855 | 0.16 | 0.709 | 1.95 | 2.02 | 0.582 | 0.42 | 0.403 | 2.13 | 2.46 | 0.280 | 0.32 | 0.534 | 2.42 | 2.65 | 0.585 |
|  |  | High-High | 2.64 | 2.08 | 0.180 | 0.87 | 0.085 | 1.71 | 2.02 | 0.033 | 0.60 | 0.254 | 2.42 | 2.46 | 0.902 | -0.57 | 0.302 | 3.78 | 2.65 | 0.014 |
| Forgotten | LTM | Low-High | 2.18 | 2.08 | 0.874 | -0.07 | 0.898 | 2.19 | 2.02 | 0.44 | 0.52 | 0.382 | 2.04 | 2.46 | 0.404 | 0.34 | 0.529 | 2.41 | 2.65 | 0.662 |
|  |  | High-High | 1.81 | 2.08 | 0.624 | 0.23 | 0.656 | 1.52 | 2.02 | 0.028 | -0.04 | 0.944 | 2.23 | 2.46 | 0.682 | -0.02 | 0.961 | 2.40 | 2.65 | 0.645 |
|  | WM | Low-High | 5.26 | 2.08 | 0.015 | 3.44 | **0.013** | 1.76 | 2.02 | 0.334 | 3.51 | **0.018** | 2.13 | 2.46 | 0.629 | 3.29 | **0.037** | 2.54 | 2.65 | 0.884 |
|  |  | High-High | 2.17 | 2.08 | 0.892 | 0.29 | 0.601 | 1.82 | 2.02 | 0.406 | -0.10 | 0.870 | 2.65 | 2.46 | 0.780 | 0.07 | 0.903 | 2.67 | 2.65 | 0.966 |

**Table 2.** Generic memory signal (mean) across relative levels of spiking at encoding and brain regions in the continuous recognition task for remembered and forgotten items. Repeated and novel distributions consisted of normalized single-neuron spike counts, with each observation representing the response of one neuron to one item. Repeated items were categorized according to the relative levels of spiking immediately before and during encoding. Primary analyses are shown for relative increases in firing (Low-High) and consistently elevated firing (High-High); results for all four firing patterns are reported in Supplementary Tables S2-S5. Retrieval demand was manipulated by lag, with long-term memory (LTM) corresponding to lags 7, 15, and 31 and working memory (WM) corresponding to lags 0, 1, and 3. ΔH-Region reflects the difference between the hippocampal repeated-novel effect and the corresponding repeated-novel effect in the comparison region. Statistical comparisons were assessed using two-sided bootstrap tests (10,000 resamples). Bolded *P* values survived Benjamini-Hochberg false-discovery-rate correction.

| Mean |  |  |  |  |  |  |  |  |  |  |  |  |  |  |  |  |  |  |  |  |
| --- | --- | --- | --- | --- | --- | --- | --- | --- | --- | --- | --- | --- | --- | --- | --- | --- | --- | --- | --- | --- |
|  |  |  | Hippocampus |  |  | Amygdala |  |  |  |  | Anterior Cingulate Cortex |  |  |  |  | Prefrontal Cortex |  |  |  |  |
| | | Firing Pattern | Repeated | Novel | P | $\Delta H-A$ | P | Repeated | Novel | P | $\Delta H-AC$ | P | Repeated | Novel | P | $\Delta H-PF$ | P | Repeated | Novel | P |
| Remembered | LTM | Low-High | -0.01 | -0.01 | 0.741 | -0.01 | 0.628 | 0.05 | 0.04 | 0.728 | -0.07 | **0.001** | 0.10 | 0.03 | **<0.001** | 0.07 | **0.004** | -0.10 | -0.02 | **<0.001** |
|  |  | High-High | 0.07 | -0.01 | **<0.001** | 0.09 | **<0.001** | 0.03 | 0.04 | 0.312 | 0.04 | **0.037** | 0.07 | 0.03 | **0.006** | 0.10 | **<0.001** | -0.04 | -0.02 | 0.199 |
|  | WM | Low-High | 0.001 | -0.01 | 0.515 | -0.04 | 0.057 | 0.09 | 0.04 | **0.001** | -0.02 | 0.376 | 0.06 | 0.03 | 0.073 | 0.03 | 0.148 | -0.04 | -0.02 | 0.182 |
|  |  | High-High | 0.09 | -0.01 | **<0.001** | 0.07 | **<0.001** | 0.07 | 0.04 | 0.014 | 0.01 | 0.762 | 0.12 | 0.03 | **<0.001** | 0.04 | 0.053 | 0.04 | -0.02 | **<0.001** |
| Forgotten | LTM | Low-High | 0.001 | -0.01 | 0.730 | -0.01 | 0.694 | 0.07 | 0.04 | 0.375 | -0.03 | 0.373 | 0.07 | 0.03 | 0.113 | -0.04 | 0.327 | 0.02 | -0.02 | 0.097 |
|  |  | High-High | 0.03 | -0.01 | 0.070 | 0.03 | 0.297 | 0.05 | 0.04 | 0.796 | 0.04 | 0.239 | 0.03 | 0.03 | 0.970 | 0.06 | 0.110 | -0.04 | -0.02 | 0.567 |
|  | WM | Low-High | 0.02 | -0.01 | 0.443 | 0.01 | 0.848 | 0.06 | 0.04 | 0.643 | 0.07 | 0.191 | -0.01 | 0.03 | 0.278 | 0.02 | 0.720 | -0.02 | -0.02 | 0.852 |
|  |  | High-High | 0.09 | -0.01 | **<0.001** | 0.05 | 0.188 | 0.09 | 0.04 | 0.107 | 0.12 | **0.009** | 0.01 | 0.03 | 0.564 | 0.17 | **<0.001** | -0.09 | -0.02 | 0.039 |

**Figure 1.**
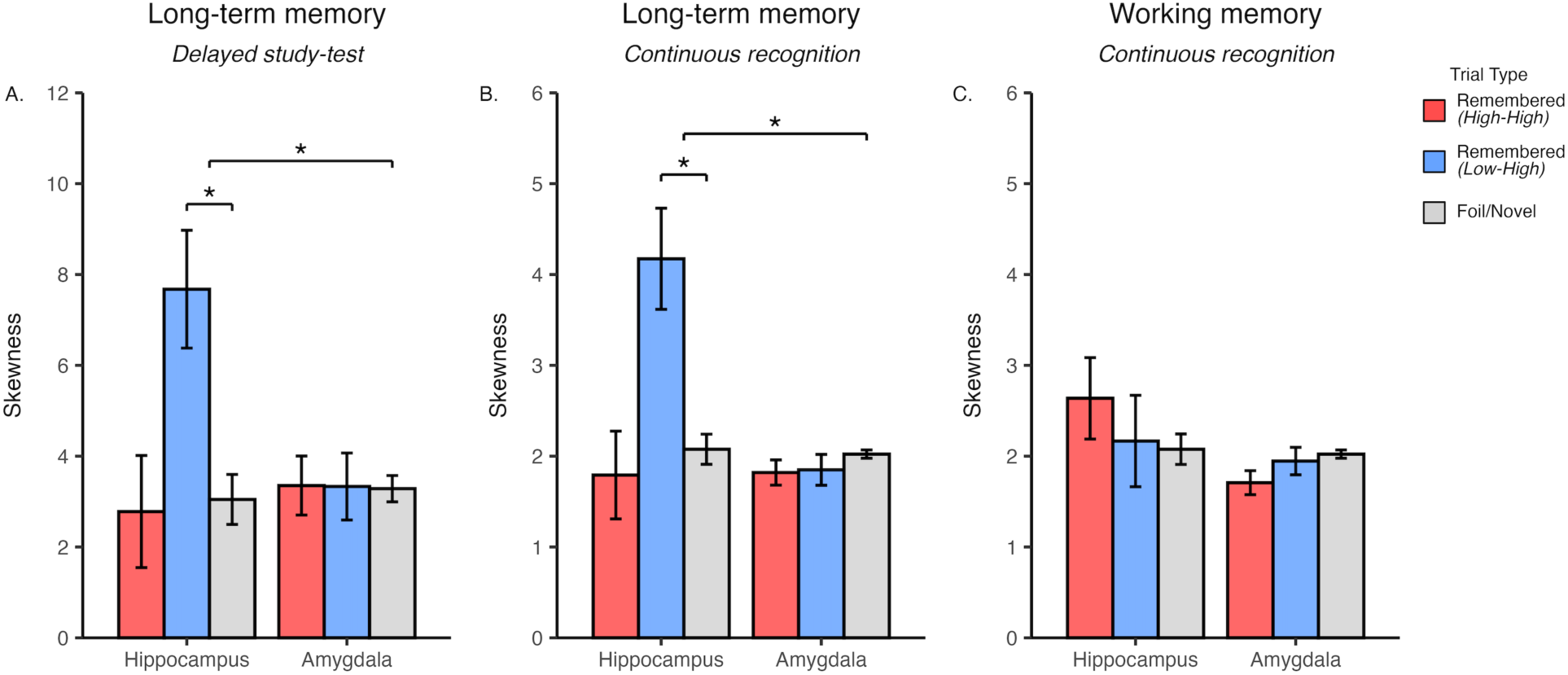
Greater skewness of the repeated than novel spike-count distributions was present only in the hippocampus during long-term memory retrieval of remembered items with relative increases in spiking at encoding. Skewness of normalized single-neuron spike-count distributions during recognition memory. (A) Independent delayed study-test recognition. (B) Long-term memory (LTM) during continuous recognition. (C) Working memory (WM) during continuous recognition. Bars summarize the skewness of normalized single-neuron spike-count distributions, with each observation representing the response of one neuron to one item. In each dataset, remembered items were categorized according to the relative levels of spiking activity immediately before and during encoding: relative increases in firing (Low-High; blue) or consistently elevated firing (High-High; red). Remembered targets are shown in A, and remembered repeated items are shown in B and C. Foils (A) and novel items (B,C) are shown in gray. Error bars represent the standard deviation of the bootstrap distribution. Brackets denote statistically significant comparisons.

### Behavioral performance differed between working memory and long-term memory retrieval demand but not across encoding firing patterns

Participants performed above chance and below ceiling across all trials, with mean session-level discriminability of *d′* = 2.13 ± 0.11 (SEM) across all sessions. Session-level analyses showed high discriminability in both retrieval demand conditions, with greater performance during working memory (WM: d′ = 2.41 ± 0.11) than long-term memory retrieval (LTM: d′ = 1.90 ± 0.11; t(54) = 7.52, P < 0.001, two-sided paired t-test). This difference was driven primarily by reduced hit rates at longer lags (WM: 86.9% ± 2.2%; LTM: 76.0% ± 3.1%), whereas false-alarm rates remained similar across working memory and long-term memory trials (WM: 18.2% ± 2.3%; LTM: 18.0% ± 2.2%). Behavioral performance also did not differ across items grouped according to the relative change in hippocampal firing immediately before and during their first presentation (repeated-measures ANOVAs; N = 53 sessions with all four encoding conditions; hit rate: F(3,156) = 1.22, P = 0.304; false-alarm rate: F(3,156) = 1.86, P = 0.139; *d′*: F(3,156) = 1.24, P = 0.296). Differences in the reported item-specific and generic neural signals across these groups were not readily explained by differences in behavioral performance.

### The hippocampal item-specific memory signal emerged selectively during long-term memory retrieval for remembered items associated with relative increases in spiking activity at encoding

We first examined whether the item-specific memory signal depended on retrieval demand, the relative levels of spiking activity before and during item presentation at encoding, subsequent memory outcome, and anatomical region. The item-specific memory signal was defined as greater skewness of the repeated-compared to novel item spike-count distribution (third moment), reflecting a sparse upper tail of disproportionately strong single-neuron responses to a small number of repeated items.

The item-specific memory signal was present only for remembered items during long-term memory retrieval that were associated with a relative increase in hippocampal spiking activity at encoding (Low-High: Table 1; Fig. 1B). Skewness of the repeated-item spike-count distribution was significantly greater than that of the novel-item distribution (Repeated = 4.17, Novel = 2.08, P = 0.006, two-sided bootstrap test: B = 10,000). The corresponding amygdala comparison was not significant (Repeated = 1.85, Novel = 2.02, P = 0.263), and the hippocampal repeated-minus-novel contrast was greater than that in the amygdala (ΔH-A = 2.26, P = 0.004, two-sided bootstrap test: B = 10,000). The same selectivity of the item-specific memory signal to the hippocampus but not the amygdala, and for only remembered Low-High repeated items, was previously observed in the independent dataset ^24^ (adapted Fig. 1A). Skewness for remembered items associated with consistently elevated firing before and during encoding (High-High) did not differ between repeated and novel spike-count distributions during long-term memory retrieval (Hippocampus: Repeated = 1.79, Novel = 2.08, P = 0.505; Amygdala: Repeated = 1.82, Novel = 2.02, P = 0.138).

In the present continuous recognition dataset, the comparison of neocortical single-neuron responses to individual items did not differ in skewness between remembered Low-High and novel items (ACC: Repeated = 2.97, Novel = 2.46, P = 0.123; PFC: Repeated = 2.59, Novel = 2.65, P = 0.907), and the hippocampal effect was greater than the corresponding effects in ACC (ΔH-AC = 1.58, P = 0.040) and PFC (ΔH-PF = 2.15, P = 0.016). Remembered items associated with relative decreases (High-Low) or consistently low firing (Low-Low) at encoding also did not differ in skewness between repeated and novel distributions (Table S2; Fig. S1).

For recognition within working memory capacity, the skewness of the repeated compared to the novel item spike-count distribution did not differ within the hippocampus, amygdala, ACC, or PFC for remembered items associated with relative increases or consistently elevated firing during encoding (Table 1; Fig. 1C: Low-High and High-High). Forgotten items associated with relative increases in hippocampal encoding activity showed numerically greater skewness for repeated than novel items during working memory (Repeated = 5.26, Novel = 2.08, P = 0.015), but this effect did not survive correction for multiple comparisons (Table S2; Fig. S2).

Kurtosis analyses, a complementary measure of tail behavior compared to skewness, supported the same hippocampal selectivity for remembered Low-High items during long-term memory retrieval (Repeated = 63.94, Novel = 9.40, P = 0.003). This effect was greater than those observed in the amygdala (ΔH-A = 56.50, P = 0.003), ACC (ΔH-AC = 45.90, P = 0.014), and PFC (ΔH-PF = 57.35, P = 0.004; Fig. S1; Table S4). Standard-deviation analyses did not show the item-specific Low-High pattern, with no difference between the repeated and novel item hippocampal spike-count distributions (Repeated = 1.02, Novel = 0.99, *P* = 0.306; Fig. S1; Table S5), suggesting that differences in skewness were not simply attributable to spike-count variability. The item-specific memory signal remained selective to remembered Low-High items during long-term memory retrieval and was not observed under the other retrieval-demand or encoding-defined conditions.

### The hippocampal item-specific memory signal was concentrated within rare, high-firing single-neuron responses to individual items

Sparse coding predicts that individual memories are represented by relatively few high-firing neurons. Within the spike-count distribution, this coding pattern would consist of relatively few disproportionately large single-neuron responses to repeated but not novel items. Skewness provides a summary measure of distributional asymmetry but does not determine whether the rare, high-firing single-neuron responses contributing to that asymmetry are concentrated within the uppermost quantiles of the spike-count distribution. Repeated compared to novel spike-count differences were therefore visualized across progressively higher response quantiles, representing progressively smaller subsets of neuron-by-item responses, using quantile-difference curves (Fig. 2A-C). Solid lines represent the hippocampus, dashed lines represent the amygdala, and the shaded gray region denotes the predefined upper-tail range used to localize the item-specific memory signal.

**Figure 2.**
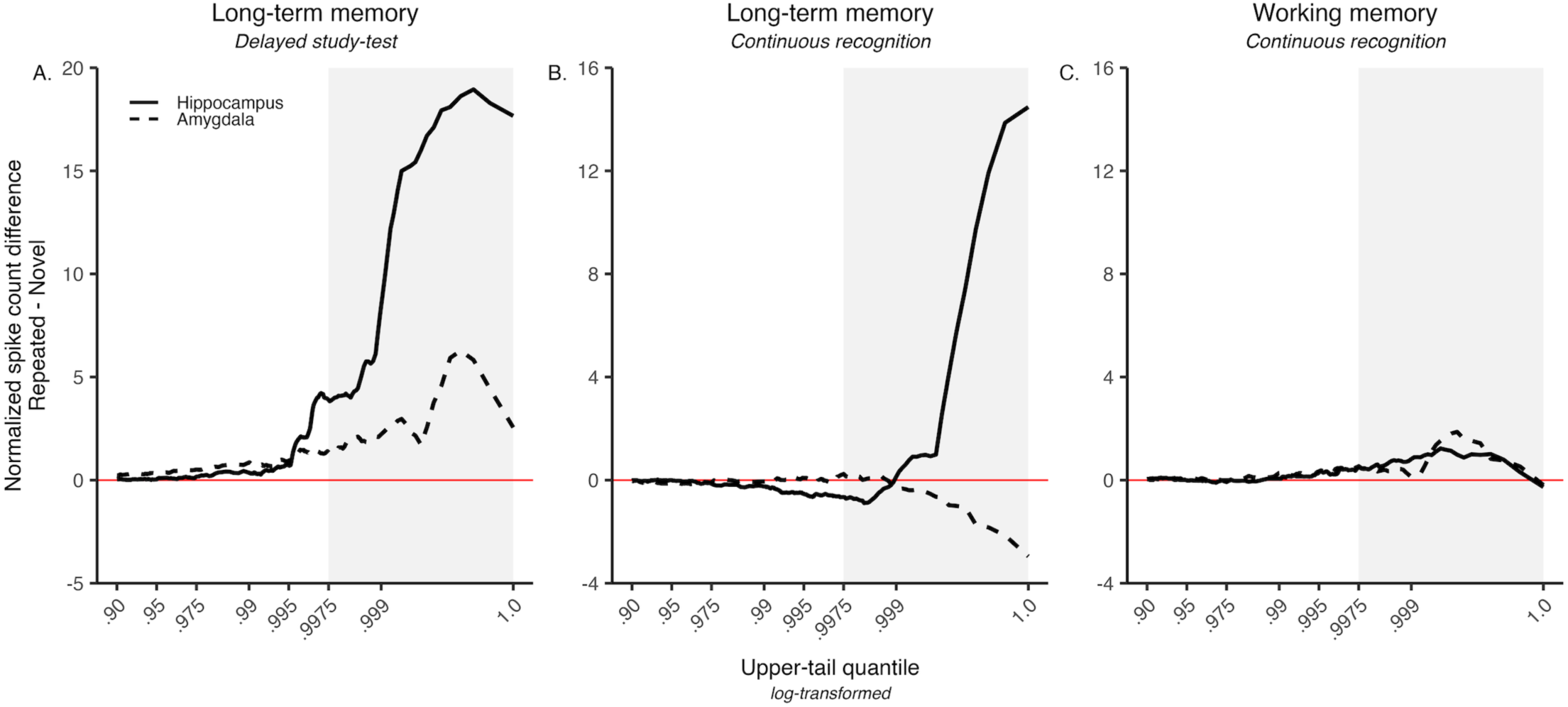
The hippocampal item-specific recognition signal was localized to rare, high-firing neuron-by-item responses in the upper tail of the spike-count distributions. Upper-tail quantile differences between normalized single-neuron spike-count distributions comparing remembered targets and foils (delayed study-test) or remembered repeated and novel items (continuous recognition). Solid lines represent the hippocampus, and dashed lines represent the amygdala. (A) Independent delayed study-test recognition. (B) Long-term memory (LTM) during continuous recognition. (C) Working memory (WM) during continuous recognition. Curves show differences in normalized single-neuron spike counts between corresponding upper-tail quantiles of the comparison distributions. As quantiles increase from left to right, progressively smaller subsets of neuron-by-item responses are represented, culminating in the highest-firing neuron-by-item responses within the upper tail of each distribution. Positive values indicate larger normalized spike counts for remembered than comparison items at each quantile (targets versus foils in A; repeated versus novel in B and C). The x-axis is logarithmically transformed to expand the upper tail of the response distributions. Shaded regions highlight the upper 0.25% of neuron-by-item responses in each distribution.

Quantile difference curves showed that the hippocampal item-specific memory signal was concentrated within rare, high-firing retrieval events. During long-term memory retrieval, the difference in spiking between repeated and novel items remained near zero across most of the quantiles and increased sharply only within the upper-most quantiles (> 0.9975) in both the delayed study-test dataset (Fig. 2A) and the continuous recognition dataset (Fig. 2B). Thus, repeated-novel differences were restricted to a small subset of disproportionately large hippocampal responses. In contrast, quantile-difference curves remained relatively flat and near zero for the hippocampus during working memory (Fig. 2C) and the amygdala showed no positive upper-tail deviation under either retrieval-demand condition (Fig. 2A-C), suggesting no evidence of disproportionately greater responses to repeated items.

### Consistently elevated firing at encoding predicted a generic recognition signal within the hippocampus and neocortex during working memory

We next tested whether the generic recognition signal also depended on retrieval demand, encoding activity, subsequent memory, and anatomical region (Fig. 3; Table 2). The generic recognition signal was present for remembered items associated with consistently elevated levels of spiking activity (High-High) immediately before and during presentation of the same item at encoding. During working memory recognition, remembered High-High repeated items showed greater mean firing than novel items in the hippocampus (Repeated = 0.09, Novel = -0.01, P < 0.001), ACC (Repeated = 0.12, Novel = 0.03, P < 0.001), and PFC (Repeated = 0.04, Novel = -0.02, P < 0.001). The amygdala also showed numerically greater repeated than novel firing (Repeated = 0.07, Novel = 0.04, P = 0.014) but did not pass correction for multiple comparisons. During long-term memory retrieval, differences in mean firing between remembered High-High compared to novel items remained significant in the hippocampus (Repeated = 0.07, Novel = -0.01, P < 0.001) and ACC (Repeated = 0.07, Novel = 0.03, P = 0.006), but not in the amygdala or PFC (Table 2). During working memory, the generic signal extended beyond the medial temporal lobe into frontal neocortex but was observed only in the hippocampus and ACC during long-term memory retrieval.

**Figure 3.**
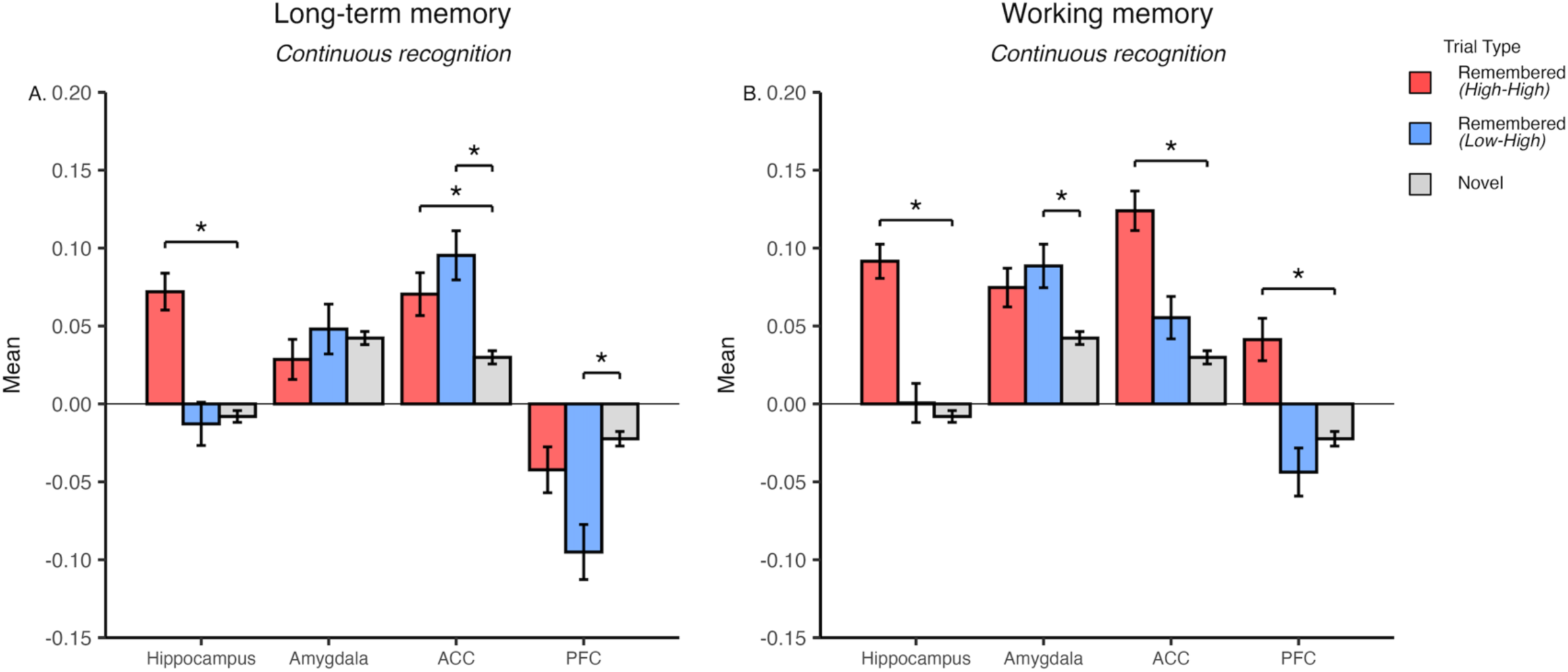
Greater mean firing of the repeated than novel spike-count distributions was present in the hippocampus and neocortex across retrieval demands for remembered items with consistently elevated spiking at encoding. Mean of normalized single-neuron spike-count distributions during continuous recognition memory. (A) Long-term memory (LTM). (B) Working memory (WM). Bars summarize the mean of normalized single-neuron spike-count distributions, with each observation representing the response of one neuron to one item. Remembered repeated items were categorized according to the relative levels of spiking activity immediately before and during encoding: relative increases in firing (Low-High, blue) or consistently elevated firing (High-High, red). Novel items are shown in gray. Error bars represent the standard deviation of the bootstrap distribution. Brackets denote statistically significant comparisons that survived Benjamini-Hochberg false-discovery-rate correction.

Within the hippocampus, remembered Low-High items did not express the same generic recognition signal. No difference in mean firing was present for long-term memory retrieval (Repeated = -0.01, Novel = -0.01, P = 0.741) or working memory (Repeated = 0.001, Novel = -0.01, P = 0.515), despite the same condition expressing the hippocampal item-specific recognition signal (Fig. 1; Table 1). Mean firing differences during working memory were also absent in the ACC and PFC. Only the amygdala showed greater mean firing during working memory for repeated Low-High items than novel items (Repeated = 0.09, Novel = 0.04, P = 0.001), but not during long-term memory retrieval (Repeated = 0.05, Novel = 0.04, P = 0.728). In the neocortex during long-term memory retrieval, remembered Low-High items showed greater mean firing than novel items in the ACC (Repeated = 0.10, Novel = 0.03, P < 0.001) and lower mean firing than novel items in the PFC (Repeated = −0.10, Novel = −0.02, P < 0.001).

### Retrieval demand and encoding activity dissociated hippocampal expression of item-specific and generic recognition signals

We next contrasted the item-specific (skewness) and generic (mean) signals established in Figures 1 and 3 directly between long-term and working memory, as a function of encoding firing state (Fig. 4; Table 3). If retrieval demand selectively influences sparse episodic retrieval, the difference in repeated-versus-novel skewness between Low-High and High-High items should be larger during long-term memory than working memory, with no corresponding change for generic mean firing.

**Figure 4.**
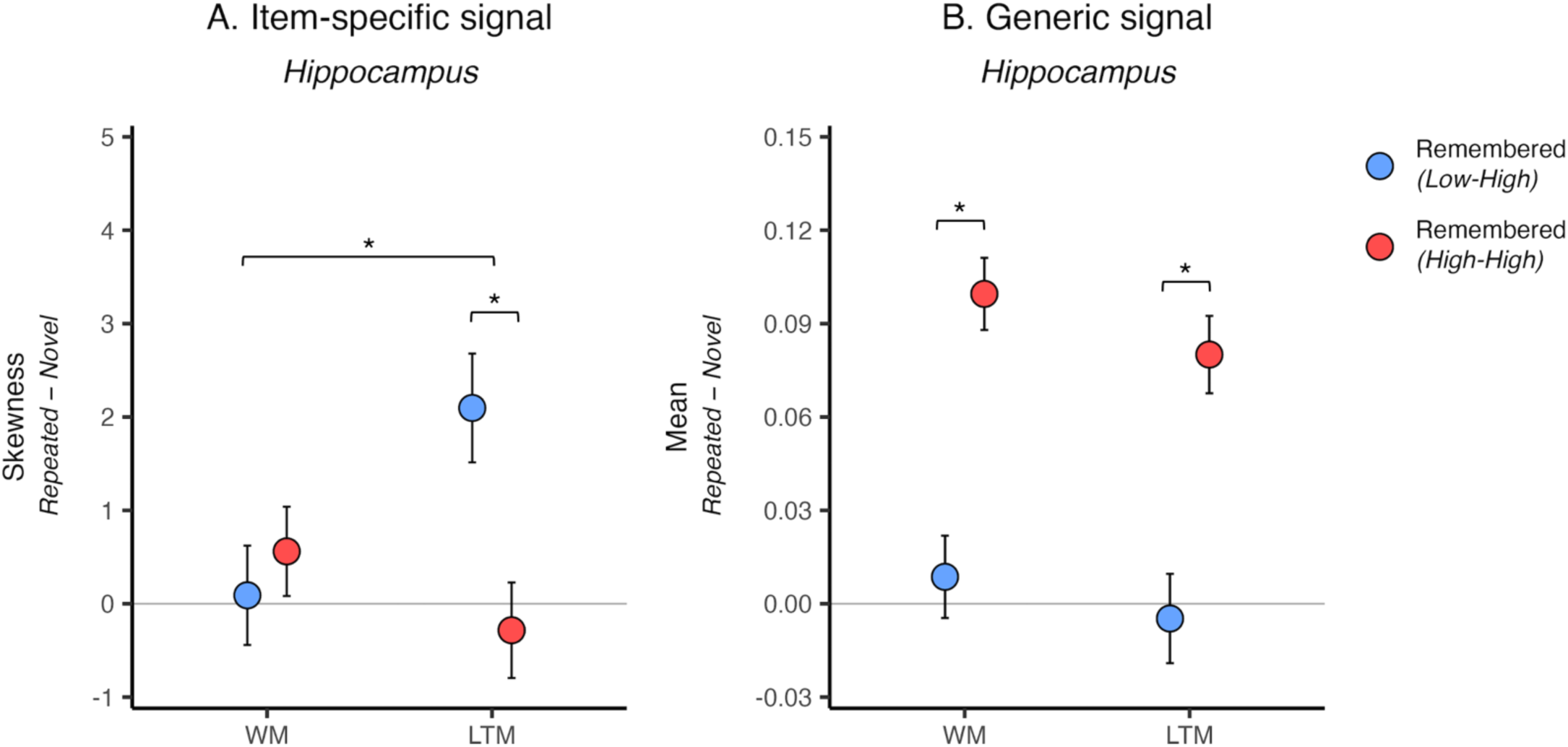
Relative spiking at encoding distinguished hippocampal item-specific and generic recognition signals across retrieval demand. Direct comparisons within the hippocampus contrasted the item-specific and generic recognition signals across encoding-associated firing patterns and retrieval-demand conditions. (A) Item-specific signal. (B) Generic signal. Points represent hippocampal remembered repeated-novel difference scores in skewness (A) or mean (B) for each encoding-associated firing pattern within each retrieval-demand condition. The underlying distributions consisted of normalized single-neuron spike counts, with each observation representing the response of one neuron to one item. Remembered repeated items were categorized according to the relative levels of spiking activity immediately before and during encoding: relative increases in firing (Low-High, blue) or consistently elevated firing (High-High, red). Error bars represent the standard deviation of the bootstrap distribution. Brackets within each retrieval-demand condition reflect direct comparisons between the Low-High and High-High encoding-associated firing patterns. Brackets across retrieval-demand conditions reflect the LTM-WM difference-of-differences comparison. Repeated-novel difference scores for each point are reported in Tables 1 and 2. Direct comparisons between the Low-High and High-High conditions and the LTM-WM difference-of-differences are reported in Table 3.

**Table 3.** Item-specific (skewness) and generic (mean) memory signals compared across encoding firing patterns and retrieval demands. Each value summarizes a distribution of normalized single-neuron spike counts, with each observation representing the response of one neuron to one item. Within the long-term memory (LTM) and working memory (WM) columns, Low-High (LH) and High-High (HH) values show the repeated-minus-novel difference in skewness or mean for remembered and forgotten items (see Tables 1–2 for full reporting). The Low-High minus High-High contrast (Δ) tests whether each memory signal differed by firing pattern at encoding within a given retrieval condition. The LTM–WM column reports the difference of differences (ΔΔ), calculated as LTM(LH − HH) − WM(LH − HH), which tests whether this encoding-related difference depended on retrieval demand. Statistical comparisons were assessed using two-sided bootstrap tests (10,000 resamples). Bolded *P* values survived Benjamini–Hochberg false-discovery-rate correction. LH–HH contrasts were corrected within each region of interest and statistic across four tests defined by subsequent-memory outcome and retrieval-demand condition. LTM–WM difference-of-differences contrasts were corrected within each region of interest and statistic across the two subsequent-memory outcomes.

|  |  |  | Long-term memory |  |  |  | Working memory |  |  |  | LTM–WM |  |
| --- | --- | --- | --- | --- | --- | --- | --- | --- | --- | --- | --- | --- |
| Moment | Subsequent Memory | Region | Low-High | High-High | Low-High – High-High |  | Low-High | High-High | Low-High – High-High |  | LH–HH |  |
| | | | | | $\Delta$ | $P$ | | | $\Delta$ | $P$ | $\Delta\Delta$ | $P$ |
| Skewness | Remembered | Hippocampus | 2.09 | -0.29 | 2.38 | **0.005** | 0.09 | 0.56 | -0.47 | 0.333 | 2.85 | **0.009** |
|  |  | Amygdala | -0.17 | -0.20 | 0.03 | 0.868 | -0.08 | -0.31 | 0.23 | 0.197 | -0.21 | 0.438 |
|  |  | ACC | 0.51 | 0.28 | 0.23 | 0.571 | -0.33 | -0.04 | -0.29 | 0.450 | 0.53 | 0.369 |
|  |  | PFC | -0.06 | -0.31 | 0.25 | 0.537 | -0.23 | 1.13 | -1.36 | 0.029 | 1.61 | 0.062 |
|  | Forgotten | Hippocampus | 0.10 | -0.27 | 0.37 | 0.426 | 3.19 | 0.09 | 3.10 | 0.037 | -2.73 | 0.094 |
|  |  | Amygdala | 0.17 | -0.50 | 0.67 | 0.054 | -0.27 | -0.21 | -0.06 | 0.852 | 0.73 | 0.152 |
|  |  | ACC | -0.42 | -0.23 | -0.19 | 0.722 | -0.33 | 0.19 | -0.52 | 0.400 | 0.33 | 0.726 |
|  |  | PFC | -0.23 | -0.24 | 0.01 | 0.981 | -0.10 | 0.03 | -0.13 | 0.795 | 0.14 | 0.855 |
| Mean | Remembered | Hippocampus | -0.005 | 0.08 | -0.09 | **<0.001** | 0.009 | 0.10 | -0.09 | **<0.001** | 0.006 | 0.800 |
|  |  | Amygdala | 0.01 | -0.01 | 0.02 | 0.335 | 0.05 | 0.03 | 0.02 | 0.449 | 0.006 | 0.835 |
|  |  | ACC | 0.07 | 0.04 | 0.03 | 0.231 | 0.03 | 0.09 | -0.06 | **<0.001** | 0.09 | **<0.001** |
|  |  | PFC | -0.07 | -0.02 | -0.05 | **0.020** | -0.02 | 0.06 | -0.08 | **<0.001** | 0.03 | 0.299 |
|  | Forgotten | Hippocampus | 0.009 | 0.04 | -0.03 | 0.346 | 0.02 | 0.10 | -0.08 | 0.079 | 0.05 | 0.403 |
|  |  | Amygdala | 0.02 | 0.006 | 0.01 | 0.619 | 0.02 | 0.05 | -0.03 | 0.478 | 0.04 | 0.386 |
|  |  | ACC | 0.04 | -0.001 | 0.04 | 0.238 | -0.04 | -0.02 | -0.02 | 0.662 | 0.06 | 0.306 |
|  |  | PFC | 0.05 | -0.02 | 0.07 | 0.107 | 0.007 | -0.07 | 0.08 | 0.127 | -0.01 | 0.774 |

In the hippocampus, repeated-versus-novel skewness for remembered Low-High items was greater for long-term compared to working memory (LTM: 2.09; WM: 0.09; LTM − WM = 2.00), whereas skewness for High-High items was lower (LTM: −0.29; WM: 0.56; LTM − WM = −0.85). As a result, Low-High skewness was greater than High-High skewness for long-term memory (Δ = 2.38, P = 0.005) but not working memory (Δ = −0.47, P = 0.333), and this difference was significantly larger during long-term than working memory (ΔΔ = 2.85, P = 0.009). The amygdala, ACC and PFC showed no such modulation by retrieval demand (Table 3).

In contrast, hippocampal repeated-versus-novel mean firing was comparable across retrieval demands for both Low-High (LTM: −0.005; WM: 0.009; LTM − WM = −0.014) and High-High items (LTM: 0.08; WM: 0.10; LTM − WM = −0.02). Mean firing was greater for High-High than Low-High items during both long-term (Δ = −0.09, P < 0.001) and working memory (Δ = −0.09, P < 0.001), and this effect did not depend on retrieval demand (ΔΔ = 0.006, P = 0.800). For forgotten items, the hippocampal High-High generic signal was present during working memory (Repeated = 0.09, Novel = −0.01, P < 0.001) but not long-term memory (Repeated = 0.03, Novel = −0.01, P = 0.070), and no forgotten-item contrast in Table 3 survived correction (Tables 2–3).

### Encoding activity dissociated neocortical expression of the generic recognition signal during working memory

We next asked whether the encoding-dependent generic signal extended beyond the hippocampus (Fig. 5; Table 3). In the ACC, repeated-versus-novel mean firing for remembered Low-High items was greater for long-term than working memory (LTM: 0.07; WM: 0.03; LTM − WM = 0.04), whereas mean firing for High-High items was lower (LTM: 0.04; WM: 0.09; LTM − WM = −0.05). As a result, mean firing was greater for High-High than Low-High items during working memory (Δ = −0.06, P < 0.001) but not long-term memory (Δ = 0.03, P = 0.231), and this difference was significantly larger during working than long-term memory (ΔΔ = 0.09, P < 0.001).

**Figure 5.**
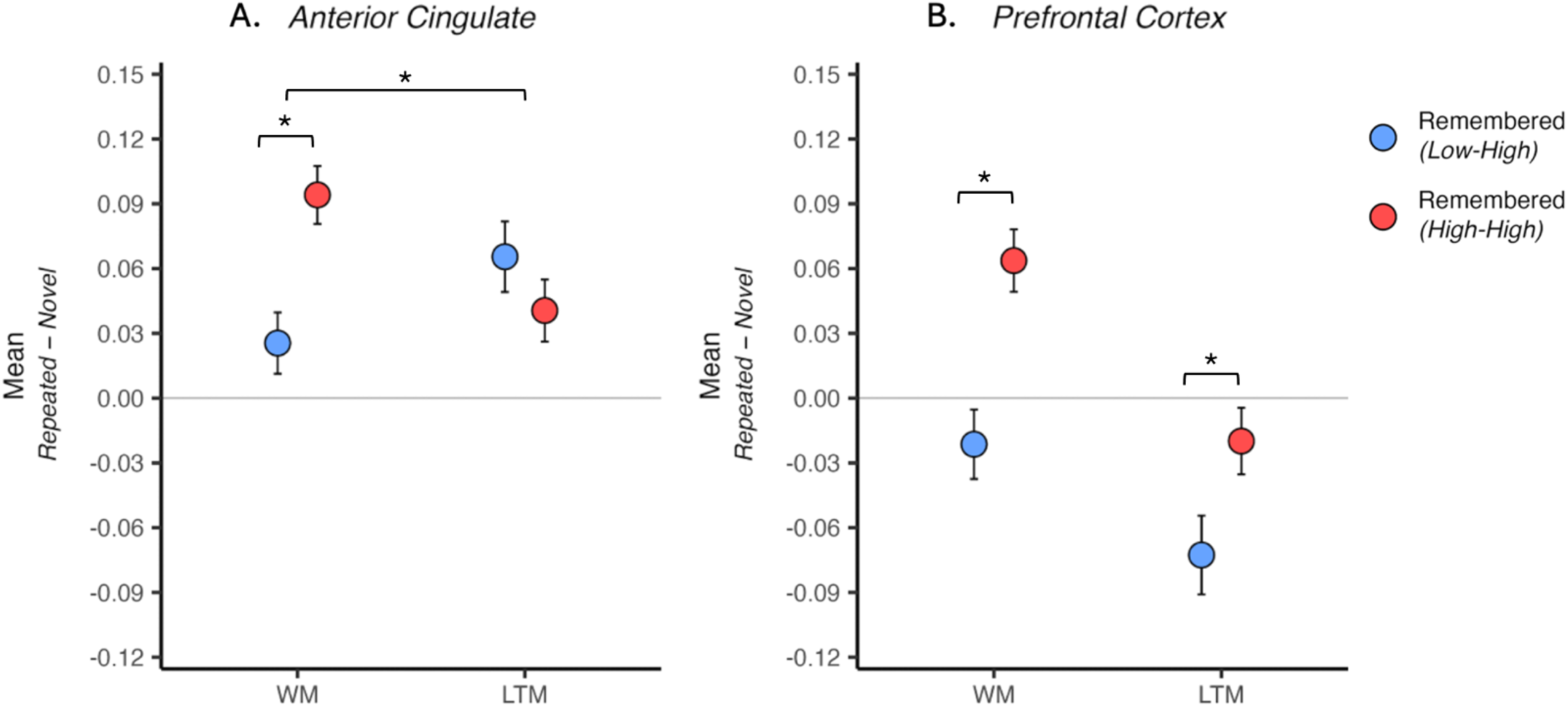
The neocortical generic recognition signal during working memory was greater for items with consistently elevated than for items with relative increases in spiking at encoding. Direct comparisons within the neocortex contrasted the generic recognition signal across encoding-associated firing patterns and retrieval-demand conditions. (A) Anterior cingulate cortex (ACC). (B) Prefrontal cortex (PFC). Points represent remembered repeated-novel difference scores in the mean of normalized single-neuron spike-count distributions for each encoding-associated firing pattern within each retrieval-demand condition. The underlying distributions consisted of normalized single-neuron spike counts, with each observation representing the response of one neuron to one item. Remembered repeated items were categorized according to the relative levels of spiking activity immediately before and during encoding: relative increases in firing (Low-High, blue) or consistently elevated firing (High-High, red). Error bars represent the standard deviation of the bootstrap distribution. Brackets within each retrieval-demand condition reflect direct comparisons between the Low-High and High-High encoding-associated firing patterns. Brackets across retrieval-demand conditions reflect the LTM-WM difference-of-differences comparison. Repeated-novel difference scores for each point are reported in Table 2. Direct comparisons between the Low-High and High-High conditions and the LTM-WM difference-of-differences are reported in Table 3.

In the PFC, repeated-versus-novel mean firing was lower during long-term than working memory for both Low-High (LTM: −0.07; WM: −0.02; LTM − WM = −0.05) and High-High items (LTM: −0.02; WM: 0.06; LTM − WM = −0.08). Mean firing was greater for High-High than Low-High items during both working (Δ = −0.08, P < 0.001) and long-term memory (Δ = −0.05, P = 0.020), with the latter reflecting lower firing for Low-High than novel items, and this effect did not depend on retrieval demand (ΔΔ = 0.03, P = 0.299).

Thus, during working memory the generic signal for High-High items was distributed across the hippocampus, ACC and PFC and was greater than the corresponding comparison for Low-High items in all three regions. This dissociation persisted during long-term memory in the hippocampus and PFC but not the ACC, the only region in which it depended on retrieval demand.

## Discussion

Recognition memory for individual words was associated with two neural signals that differed in their selectivity according to retrieval demand and encoding state. The hippocampal item-specific signal, greater positive skewness for repeated than novel spike-count distributions, was observed for remembered items under long-term-memory retrieval demands and specifically for items showing a transition from relatively low pre-encoding firing to relatively high firing during encoding (Fig. 1; Table 1). Quantile analyses localized this effect to a small number of rare, high-firing hippocampal neuron-by-item responses (Fig. 2). By contrast, the generic repeated-novel mean-firing signal was associated with consistently high firing before and during encoding and was evident in the hippocampus across both working- and long-term-memory conditions (Figs. 3-4; Tables 2-3). Its expression extended more broadly into frontal cortex during working memory (Fig. 5; Tables 2-3). Direct hippocampal contrasts within the same task and neuronal populations showed that increasing the computational demand to retrieve individual episodic representations enhanced the item-specific signal but not the generic signal, providing the clearest dissociation between the two forms of recognition activity.

These findings fit an updated Atkinson-Shiffrin framework that links short- and long-term memory to the complementary coding schemes proposed by Complementary Learning Systems (CLS) theory ^12^. In this account, recently encountered information can remain directly accessible in a transient, task-ready cortical state instantiated by overlapping neural codes. A sparse, pattern-separated hippocampal index can be formed in parallel, but retrieval through that index need not contribute measurably to performance while the cortical state remains sufficiently accessible. As subsequent input recruits the same overlapping cortical populations, the earlier cortical state becomes increasingly reconfigured and blurred ^13^. Under those conditions, a hippocampal index can provide episode-specific access by reinstating the associated cortical representation ^34,35^. Thus, long-term-memory retrieval does not replace cortical representation with hippocampal representation. Cortical coding contributes under both conditions, while hippocampal indexing becomes increasingly consequential when direct access to the earlier cortical state is insufficient. The generic signal is therefore not expected to provide a unique marker of either cortical coding or working memory. Its persistence across levels of retrieval demand is nevertheless compatible with cortical participation under both conditions. The more diagnostic result is the selective enhancement of the item-specific hippocampal signal under long-term-memory retrieval demands that are likely to rely on a hippocampal index.

CLS proposes that hippocampal circuitry supports sparse, pattern-separated codes that preserve the discriminability of individual episodes despite interference ^10,11,15^. High-resolution fMRI provides evidence for pattern separation in the human hippocampus ^14,18^, but cannot establish whether sparse coding is expressed at the level of individual neurons. The item-specific signal has properties expected of such a code. Greater repeated-novel skewness was selective to the hippocampus, and quantile-difference curves showed that the effect arose almost entirely from the upper tail of the response distribution comprising single-neuron responses to individual items, rather than from a population-wide increase in firing. Across two recognition memory datasets, the effect remained selective to remembered long-term-memory trials and to items showing a relative increase in firing at encoding despite differences in stimuli, patients, laboratories, and spike-sorting procedures ^24,32,33^.

One alternative to a sparse episodic-coding account is that the rare, high-firing hippocampal responses reflect invariant selectivity for particular concepts ^36,37^. However, the present analyses compared responses to the same item at its first and second presentation. Concept selectivity alone therefore cannot explain why the spike-count distribution was more positively skewed for repeated than novel items. These results do not establish that the item-specific signal is itself a hippocampal index, but its neuron-level properties are consistent with the sparse, item-specific hippocampal coding expected of such an index.

The dependence of the item-specific signal on a transition from a relatively low firing to high firing encoding state may point to its underlying neural mechanism. The present continuous-recognition task additionally showed that this low to high activity encoding state was not sufficient by itself: the item-specific signal became evident when recognition imposed long-term-memory retrieval demands, but not when the same encoding pattern preceded recognition within working memory capacity. This suggests that the item-specific representation formed at encoding was accessible at retrieval depending on the computational demands. Causal studies of neuronal allocation in rodents suggest a possible mechanism linking encoding state to the assignment of neurons to a memory trace. Recruitment of neurons into an engram is competitive, with intrinsic excitability ^38–40^, learning-induced changes in excitability ^41^, and pre-existing patterns of functional connectivity among neurons biasing which neurons are allocated to an episode ^42^. Competitive recruitment has also been observed in dentate gyrus ^43^, consistent with circuitry proposed to support sparse, pattern-separated coding ^16,17^. The low-to-high firing transition observed here and in our previous study ^24^ may be an extracellular correlate of allocation-related changes in neuronal state, consistent with the proposal that excitability-based allocation assigns human hippocampal neurons to individual episodes^44^, although the cellular mechanisms themselves cannot be measured with the present recordings.

The generic recognition signal showed a different relationship to encoding. It was associated with consistently high firing before and during encoding and was present in the hippocampus under both working- and long-term-memory retrieval demands. This encoding pattern is consistent with the broader subsequent-memory literature, in which elevated prestimulus and encoding-related activity predicts later remembering ^26,27^, including hippocampal spiking in the present continuous-recognition dataset ^28^. However, a subsequent-memory effect at encoding does not by itself predict that later retrieval will be expressed as a generic repeated-novel mean-firing difference. The present results therefore extend the subsequent-memory literature by linking a consistently high encoding state to later expression of a generic recognition signal that is distinct from the item-specific signal. The dissociation between low-to-high and consistently high encoding states suggests that different neural conditions at encoding can be associated with different recognition signatures at retrieval.

The presence of the generic hippocampal signal during working memory is compatible with the idea that there is not a strict double dissociation between hippocampal involvement in working- and long-term memory ^8^. A hippocampal index may be formed in parallel while information remains directly accessible in cortex, and prior studies have shown hippocampal engagement during short-delay tasks as memory load and relational demands increase ^6,7,45–49^. Unlike long-term memory, working memory does not necessarily depend on the hippocampus. Cortical representations can support some relational information while the task-ready cortical state remains sufficient, with pattern-separated, hippocampal indexing becoming more consequential as that state is degraded or competing episodes must be distinguished ^12^. The corresponding retrieval demands during continuous recognition suggest the hippocampus can flexibly engage in different coding schemes depending on the task at hand. The generic hippocampal signal was robust during working memory, whereas the item-specific signal characteristic of remembered long-term-memory trials was not. Thus, hippocampal engagement during a working memory task should not by itself be taken as evidence that recognition depends on retrieval of a pattern-separated episodic representation.

Beyond the hippocampus, the same generic signature appeared in frontal neocortex during working memory, but it did not behave uniformly across regions or retrieval conditions. For recognition of remembered items during working memory, items surrounded by consistently high firing at encoding showed reliable repeated-novel firing differences in hippocampus, ACC, and PFC. During long-term-memory retrieval, the same generic effect remained significant in hippocampus and ACC but not PFC. The High-High versus Low-High difference was clearest in ACC and PFC during working memory. In the PFC, a smaller difference was also present during long-term-memory retrieval, driven by lower firing for Low-High compared to novel items, and the interaction with retrieval demand was significant in ACC but not PFC (Fig. 5; Table 3). These regional and retrieval-demand dependencies between the hippocampus and neocortex, and within the prefrontal cortex suggest that a shared firing-rate signature need not reflect a shared computation. Medial frontal neurons more often represent task phase and memory load than stimulus identity ^50^, and also may regulate the accessibility of representations maintained elsewhere ^51,52^. Consistent with a coordinating role, hippocampal-cortical synchronization increases with working memory demand ^53,54^. Most directly, separate hippocampal populations have been reported to participate in frontal coordination and in item-selective coding ^55^. The generic signal may therefore reflect coordinated but functionally distinct computations across hippocampus and frontal cortex rather than a common mnemonic representation.

Several qualifications are important. Analyses of forgotten items were based on substantially fewer trials and should be interpreted cautiously. Importantly, the relationship between the consistently high firing encoding state and the generic signal does not imply that expression of that signal is specific to successful recognition. The generic hippocampal signal was present for forgotten as well as remembered items during working memory, which suggests it may partly reflect repetition-sensitive or statistical-learning processes within CA1 ^56,57^. The nominal item-specific effect for forgotten working memory items did not survive correction for multiple comparisons and does not warrant a strong interpretation. More generally, lag in the present task jointly increased the number of intervening items and elapsed time, so the experiment does not isolate interference from time. Nor do the recordings directly measure reconfiguration of cortical representations or hippocampally driven cortical reinstatement. The patient population, clinically determined electrode placement, and possibility of recording from epileptogenic tissue constrain generalization. Human extracellular recordings also cannot directly measure the cellular mechanisms proposed by neuronal-allocation models ^58^ or reliably resolve recordings originating from dentate gyrus and CA3. The data therefore support, but do not directly demonstrate, the specific cortical and hippocampal mechanisms proposed by CLS and the updated Atkinson-Shiffrin framework.

## Methods

### Participants

The participants consisted of 35 patients (mean age, 41 ± 2.02 years; 21 females and 14 males; all but 2 were right-handed) with medically intractable epilepsy who required implantation of depth electrodes (Ad-Tech Medical) for clinical evaluation and consideration of possible surgical resection of their seizure foci. The patients completed sessions of a continuous recognition memory task of word learning, with individual patients completing 1 to 4 sessions. Due to low recognition memory performance (*d′* < 0), 10 sessions from 6 of the 35 patients and all sessions from 1 additional patient were excluded, resulting in 34 patients and 55 total sessions included for analysis.

### Informed Consent

Patients were evaluated for depth electrode implantation at the Barrow Neurological Institute for treatment of intractable epilepsy. Recommendations for implantation were made by treating physicians solely based upon established clinical criteria ^59^. All patients provided written informed consent for research participation under protocols approved by the Institutional Review Board of St. Joseph’s Hospital and Medical Center.

### Data Availability Statement

Anonymized single-neuron data are available within an Open Science Framework repository https://osf.io/9tgmx/ ^60^.

## Intracranial Recording

### Electrode implantation and localization

Extracellular potentials corresponding to single-neuron activity were recorded from the tips of nine 38-μm-diameter microwires implanted along with a clinical depth electrode (Ad-Tech Medical Corporation, Racine, WI) used to record field potentials ^61,62^. Depth electrode implantation sites were chosen according to clinical criteria and included bilateral placement in the amygdala, anterior cingulate cortex (ACC), hippocampus, and prefrontal cortex (PFC). Extratemporal recording sites were chosen to check whether the patients’ seizures had an origin that would change recommendations for further surgery.

Target locations within each brain region were chosen by selecting a trajectory for the depth electrode which positioned the low-impedance contacts along the shaft of the depth electrode in clinically mandated areas with the tip of the depth electrode ∼3 mm superficial from the deepest part of a target structure. Electrodes were implanted stereotactically (Medtronic Stealth Station) using a 3T structural MRI and were inserted through a skull bolt with a custom frame to align the depth electrode along the chosen trajectory. The error in tip placement using this technique is estimated to be ± 3 mm based on manual inspection of the pre-operative MRI and post-operative CT and prior work ^63,64^.

### Signal acquisition and spike sorting

Extracellular potentials were digitized at 29,412 Hz with 16-bit resolution. Possible action potential events (APs) were detected using digital filtering and thresholding ^65^. Because more than one neuron may be recorded near any given electrode, APs were sorted into several clusters of similar waveform shape using the open-source clustering program KlustaKwik (Klustakwik.sf.net). After sorting, each cluster was graded as being noise, multiunit activity (MUA), or single-unit activity (SUA) based on criteria such as the waveform shape, size of the waveform relative to noise, evidence of a refractory interval, and lack of powerline interference ^66^.

## Continuous recognition memory test

### Task design

Patients were tested using a continuous recognition memory task of word learning, as previously described ^21^. Individual words were serially presented, and each word appeared twice in the experiment. The task was to judge whether each item was novel (i.e., first presentation of the word) or repeated (i.e., second presentation of the word). The words were presented either visually on a computer screen or auditorily through headphones.

Across 55 sessions (from 34 patients), there were 35 visual sessions and 20 auditory sessions. For the visual sessions, 360 words (120 one-syllable, 120 two-syllable, and 120 three-syllable words) were used. The words were taken from the Medical Research Council Psycholinguistic database ^67^. Another set of 45 one-syllable words was used as fillers that were never repeated. Three nonoverlapping word sets could be presented, and these were used for patients who volunteered for multiple sessions. Each session comprised 255 trials (240 trials in which each of the 120 words was presented twice, along with 15 filler trials).

In each trial, a word was visually displayed on a computer screen for 1,500 ms, followed by a question mark as a response prompt. Patients had up to 2,000 ms to indicate whether the word was novel or repeated. There was a jittered intertrial interval (mean = 888 ms; SD = 552 ms). The procedure for the auditory sessions was similar. Each auditory session included 615 trials wherein 300 unique words were repeated once and 15 filler words were presented once. Auditory words had a mean presentation duration of 652 ± 65 ms, and patients had up to 1,600 ms to respond. The intertrial interval was jittered (mean = 1,053 ms; SD = 53 ms).

### Lag conditions

For each pair of items, the words were randomly assigned to different lag conditions consisting of 0, 1, 3, 7, 15, or 31 intervening items, and each item pair was equally likely to appear in each condition. Items assigned a lag of 0 were presented twice in a row with no other words in between the first presentation and second presentation. For items with lags greater than 0, the lag number corresponds to the number of intervening words presented in between the first and second presentation.

### Behavioral analyses

Behavioral analyses were conducted at the session level (n = 55) and each session was treated as an independent observational unit. Although some patients completed more than one session, the task was conducted on different days, and it is typically assumed that different neurons were recorded because depth electrodes shift slightly as patients move around. Discriminability (*d′*) was computed using the log-linear correction for infinite values ^68^. Hit-rate, false alarm rate, and discriminability were calculated across the entire session, and separately for working memory and long-term memory retrieval demand. Behavioral differences between working memory and long-term memory were evaluated using two-sided paired t-tests. Hit rates, false-alarm rates, and discriminability were also compared across the four encoding-defined firing conditions using repeated-measures ANOVAs.

### Single-neuron responses

Single-neuron responses were quantified using spike counts measured before and after stimulus presentation. Pre-stimulus spike counts were computed within a baseline window extending from 1000 ms to 200 ms before stimulus onset. Post-stimulus spike counts were computed within a retrieval-response window extending from 200 ms to 1000 ms after stimulus onset. These windows were selected to minimize contamination from stimulus-onset transient firing while capturing sustained retrieval-related firing activity.

To account for differences in baseline firing rates across neurons, post-stimulus spike counts were normalized separately for each neuron. For neuron j and item i, the normalized response was calculated as:

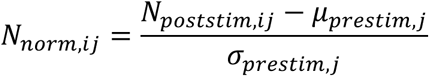

where *N_poststim_*_,*i*j_ denotes the post-stimulus spike count for neuron *j* during presentation of item *i*, and *μ*_prestim,j_ and *σ*_prestim,j_ denote the mean and standard deviation of pre-stimulus spike counts for neuron j across all trials within the session. Positive values therefore indicate greater-than-baseline firing, whereas negative values indicate lower-than-baseline firing. All distributional analyses were performed on these normalized spike-count values.

### Unit of analysis

The primary unit of analysis was the neuron-by-item observation. Each observation represented the normalized spike count for a single neuron during presentation of a single item. Following within-neuron normalization, neuron-by-item observations were pooled within each analysis partition to construct the repeated-item and novel-item spike-count distributions.

Across all sessions, recordings included 1,391 identified single units and 10,172 item presentations, yielding 485,120 neuron-by-item observations. Regional coverage included the hippocampus (396 neurons; 143,703 neuron-by-item observations), amygdala (375 neurons; 127,363 neuron-by-item observations), anterior cingulate cortex (ACC; 344 neurons; 121,961 neuron-by-item observations), and prefrontal cortex (PFC; 276 neurons; 92,093 neuron-by-item observations). Statistical analyses were conducted on pooled neuron-by-item observations rather than subject-level summary statistics, following previous distributional analyses of human single-neuron recognition memory ^21–23^. The subset of neuron-by-item observations contributing to each analysis after partitioning by retrieval demand, encoding-defined firing condition, subsequent memory outcome, and anatomical region is summarized in Table S1.

### Retrieval demand conditions (lag-defined)

Repeated words were categorized based on lag, defined as the number of intervening items between the first and second presentation of a word. Short lags (0, 1, 3) were classified as working memory (WM), and long lags (7, 15, 31) were classified as long-term memory (LTM), based on established capacity limits of working memory. During recognition within working memory capacity limits, the initial presentation is likely still active within overlapping cortical representations, whereas at long lags, successful recognition relies on retrieval from interference-resistant item-specific representations from long-term memory.

### Relative spiking activity at encoding and subsequent memory conditions (item-defined)

Repeated items were categorized into one of four encoding conditions based on the relative spiking activity elicited by the item at encoding: (1) relative increase (Low-High), (2) consistently elevated (High-High), (3) relative decrease (High-Low), and (4) consistently low (Low-Low). Within each region and session, items were partitioned using median splits applied separately to pre-stimulus onset (-1000 to -200 ms) and post-stimulus onset (200 to 1000 ms) spike counts to reflect relative changes in firing rate at encoding. Median splits were computed within session and region to avoid bias from between-session differences in firing rate. These categories are relative changes in regional firing associated with individual items and are interpreted as proxies for encoding-related neural dynamics that bias assignment of neurons to a memory trace. Repeated items were further categorized by subsequent memory outcome during recognition as remembered (hit) or forgotten (miss). Thus, repeated items were assigned labels based on the conjunction of retrieval-demand, spiking activity at encoding, and subsequent-memory outcome.

## Generic and item-specific memory signals

### Distributional analysis approach

Repeated-item and novel-item retrieval-response distributions were constructed from pooled neuron-by-item observations. Each observation corresponded to the within-session normalized spike-count of a single-neuron in response to a single item presentation. The repeated-novel difference between each of the four distributional moments (mean, standard deviation, skewness, and kurtosis) was compared separately within each brain region (hippocampus, amygdala, ACC, and PFC), and for each subset of items partitioned by retrieval-demand (working memory and long-term memory), relative spiking activity at item encoding (Low-High, High-High, High-Low, and Low-Low), and subsequent-memory outcome (remembered and forgotten). The corresponding novel-item distribution was generated within each brain region and served as the common comparison for each repeated-item distribution. Planned contrasts also directly compared the repeated-novel differences between the hippocampus and each comparison region (amygdala, ACC, and PFC), between the Low-High and High-High encoding firing patterns, and between retrieval-demand differences.

The comparisons between both normalized spike-count distributions were designed to distinguish two types of memory signals that support different aspects of recognition. For each repeated-item and novel-item distribution, the first four moments were calculated: mean, standard deviation, skewness, and kurtosis. A generic memory signal was defined as increased mean firing across many neuron-by-item observations, producing a shift in the central tendency of the repeated-item distribution relative to the corresponding novel-item distribution. In contrast, sparse-coding models predict that item-specific retrieval should be concentrated within a relatively small subset of high firing neuron-by-repeated item responses while leaving most neuron-by-item observations across both repeated and novel distributions largely similar. An item-specific recognition signal was defined as greater skewness of the repeated-item distribution relative to the novel-item distribution, reflecting rare high-firing neuron-by-item responses. Kurtosis was examined as a complementary measure of heavy-tail structure but was not interpreted as primary evidence for the item-specific recognition signal.

## Statistical comparisons

For each moment (mean, standard deviation, skewness, and kurtosis), the repeated-item and novel-item spike-count distributions were compared within each brain region. Statistical significance was assessed using bootstrap resampling (*n* = 10,000). Within each brain region, individual repeated-item and novel-item observations were sampled with replacement from the pooled distribution of normalized neuron-by-item spike counts while preserving the original sample sizes. The repeated-novel difference for each moment was calculated from the repeated-item and novel-item bootstrap samples. The resulting distribution of repeated-novel differences formed the null distribution. Two-sided P values were calculated as the proportion of absolute resampled differences greater than or equal to the absolute observed difference. Error bars represent the standard deviation of the null distribution and reflect the expected variability of the repeated-novel difference.

### Within-region repeated-versus-novel contrast

Within-region retrieval effects were quantified as the difference between repeated-item and novel-item normalized spike count distributions consisting of all neuron-by-item spike counts across all sessions:

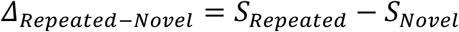

where *S* denotes the moment of interest (mean, standard deviation, skewness, kurtosis). Positive values reflect larger values for repeated-item than novel-item moment, whereas negative values reflect larger values for novel-item moment.

### Between-region interaction contrast

To assess anatomical specificity, repeated-versus-novel effects observed in the hippocampus were compared with the corresponding effects observed in each comparison region. For a given moment, the hippocampus-versus-region contrast was defined as:

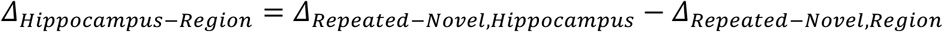

Each comparison region was tested separately (amygdala, anterior cingulate cortex, and prefrontal cortex). Positive values indicate larger repeated-novel differences in the hippocampus than in the comparison region, whereas values near zero indicate comparable repeated-novel differences across regions.

### Relative spiking activity at encoding contrast

Retrieval effects were compared according to relative levels of spiking activity immediately before and during encoding. Items that elicited relative increases in spiking activity before and during item presentation at encoding (Low-High) were compared to items exhibiting consistently elevated spiking activity at encoding (High-High). For a given moment, the relative spiking activity contrast was defined as:

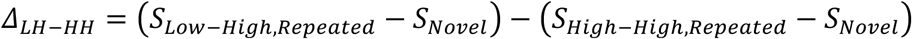

where *S_Lo_*_w–*Hi*gℎ,*Repeated*_ and *S_Hi_*_gℎ–*Hi*gℎ,*Repeated*_ denote the statistical moment of interest (mean or skewness) for the repeated-item distributions in the Low-High and High-High conditions, respectively, and *S_Novel_* denotes the statistical moment for the common novel-item distribution used for both contrasts. Positive values indicate larger repeated-novel differences for Low-High than High-High items, whereas negative values indicate larger repeated-novel differences for High-High than Low-High items. These contrasts were calculated separately within each brain region, retrieval-demand condition, moment, and subsequent-memory outcome.

### Retrieval-demand by relative spiking activity at encoding contrast

Recognition effects were compared between long-term memory (LTM) and working memory (WM) retrieval demands according to relative levels of spiking activity immediately before and during item presentation at encoding. For a given moment, the retrieval-demand contrast was defined as:

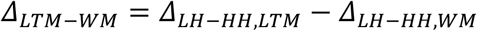

where *Δ_LH_*_–*HH*,*LTM*_ and *Δ_LH_*_–*HH*,W*M*_ denote the Low-High minus High-High, repeated-novel contrasts during long-term memory and working memory, respectively. Positive values indicate larger differences between the Low-High and High-High repeated-novel contrasts during long-term memory than working memory, whereas negative values indicate larger differences between the Low-High and High-High repeated-novel contrasts during working memory than long-term memory. Each contrast served as the inferential test of whether retrieval demand modulated the Low-High versus High-High difference for an individual region. Differences between long-term memory and working memory within a single firing pattern were not tested separately and are reported descriptively to characterize the pattern underlying a significant contrast.

### Distributional structure of the item-specific memory signal

The distributional structure underlying the item-specific recognition signal (skewness) was examined to determine whether differences between remembered items and their corresponding comparison items were concentrated within rare, high-firing neuron-by-item observations or extended more broadly across the upper tail of the spike-count distributions.

The upper-tail structure was visualized using repeated-minus-novel quantile-difference curves for remembered Low-High repeated items, which expressed the item-specific recognition signal. Empirical quantiles were computed separately for the normalized repeated-item and novel-item spike-count distributions within the hippocampus and the amygdala, which served as a medial temporal lobe comparison region. At each quantile probability *p*, the corresponding novel-item quantile value was subtracted from the repeated-item quantile value: *Q*_(*Repeated*)(*p*)_ − *Q*_(*Novel*)(*p*)_ where *Q*_(*Repeated*)(*p*)_ and *Q*_(*Novel*)(*p*)_ denote the repeated-item and novel-item spike-count quantiles, respectively. Each point represents the repeated-minus-novel difference at the corresponding quantile of the two distributions. Positive values reflect larger responses for repeated than novel items within that quantile and negative values reflect larger responses for novel than repeated items within that quantile.

The same quantile-difference analysis was independently applied to the delayed study-test recognition dataset to verify the generalizability of the item-specific recognition signal. In that dataset, remembered targets (items presented during the study phase and correctly recognized during the subsequent test phase) were compared with foils (new items presented only during the test phase).

Quantile-difference curves were displayed on a logarithmic scale to increase resolution of rare high-firing single-neuron retrieval events. Interpretation focused *a priori* on the upper 0.25% of the neuron-by-item response distribution (*p* = 0.9975-0.9999). This quantile range was selected based on prior human single-neuron recognition-memory studies showing that hippocampal repeated-minus-novel distributional differences are most prominent within this upper-tail region ^21,24^.

### Multiple comparisons

Benjamini-Hochberg false-discovery-rate correction (q < 0.05) was applied using multiple-comparison families defined by the primary inferential objective of each analysis. Repeated-novel comparisons were performed separately for four encoding-associated firing patterns (Low-High, High-High, High-Low, Low-Low), two subsequent-memory outcomes (remembered, forgotten), and four distributional statistics (mean, standard deviation, skewness, kurtosis). False-discovery-rate correction was applied within each region of interest across these 32 comparisons, separately for working memory (WM), long-term memory (LTM), and delayed study-test analyses. Hippocampus-versus-region contrasts are reported for all conditions and were corrected across the three comparison regions (amygdala, anterior cingulate cortex, and prefrontal cortex). Planned Low-High versus High-High contrasts were corrected within each region of interest and statistic across four comparisons defined by retrieval-demand condition (working memory, long-term memory) and subsequent-memory outcome (remembered, forgotten). Planned difference-of-differences contrasts were corrected within each region of interest and statistic across the two subsequent-memory outcomes.

## Author contributions

C.W.T. contributed data analysis, data interpretation, and manuscript writing. J.T.W. supervised the project and contributed to all aspects of the manuscript. P.N.S. contributed data curation and statistical guidance at all stages of the analysis and manuscript writing. All the authors discussed the results at all stages of the project.

