## Supplemental Material for "Encoding activity predicts distinct recognition signals during working and long-term memory in the human hippocampus"



**Author contributions:**

C.W.T. contributed data analysis, data interpretation, and manuscript writing. J.T.W. supervised the project and contributed to all aspects of the manuscript. P.N.S. contributed data curation and statistical guidance at all stages of the analysis and manuscript writing. All the authors discussed the results at all stages of the project.

**Competing interests:**

C.W.T., J.T.W, and P.N.S. have no competing interests.

| | | Encoding | Remembered | | | | Forgotten | | | |
|---|---|---|---|---|---|---|---|---|---|---|
| | | Novel | Low-High | High-High | Low-Low | High-Low | Low-High | High-High | Low-Low | High-Low |
| LTM | **Hippocampus** | | | | | | | | | |
| | # Items | 4,281 | 636 | 878 | 858 | 651 | 188 | 239 | 266 | 158 |
| | # Neuron-by-item pairs | 34,017 | 5,312 | 7,159 | 6,588 | 5,262 | 1,557 | 2,173 | 2,529 | 1,323 |
| | # Upper-tail neuron-by-item pairs | 87 | 7 | 15 | 20 | 9 | 5 | 4 | 6 | 1 |
| | # Upper-tail unique neurons | 68 | 6 | 15 | 19 | 9 | 4 | 3 | 6 | 1 |
| | **Amygdala** | | | | | | | | | |
| | # Items | 3,640 | 531 | 726 | 729 | 528 | 167 | 190 | 249 | 151 |
| | # Neuron-by-item pairs | 30,538 | 4,242 | 6,469 | 5,966 | 4,228 | 1,731 | 1,955 | 2,577 | 1,485 |
| | # Upper-tail neuron-by-item pairs | 81 | 12 | 13 | 15 | 12 | 9 | 1 | 8 | 5 |
| | # Upper-tail unique neurons | 58 | 11 | 13 | 13 | 10 | 8 | 1 | 7 | 5 |
| | **Anterior Cingulate Cortex** | | | | | | | | | |
| | # Items | 3,757 | 542 | 763 | 786 | 561 | 193 | 215 | 253 | 158 |
| | # Neuron-by-item pairs | 29,381 | 4,414 | 5,811 | 5,857 | 4,636 | 1,499 | 1,587 | 1,934 | 1,176 |
| | # Upper-tail neuron-by-item pairs | 78 | 14 | 15 | 4 | 6 | 5 | 5 | 6 | 2 |
| | # Upper-tail unique neurons | 58 | 14 | 15 | 4 | 6 | 4 | 5 | 6 | 2 |
| | **Prefrontal Cortex** | | | | | | | | | |
| | # Items | 3,166 | 462 | 685 | 639 | 465 | 152 | 179 | 183 | 124 |
| | # Neuron-by-item pairs | 21,821 | 3,100 | 4,474 | 4,124 | 3,153 | 1,305 | 1,351 | 1,586 | 1,000 |
| | # Upper-tail neuron-by-item pairs | 49 | 5 | 7 | 6 | 13 | 5 | 2 | 5 | 2 |
| | # Upper-tail unique neurons | 33 | 5 | 5 | 6 | 13 | 4 | 2 | 4 | 2 |
| WM | **Hippocampus** | | | | | | | | | |
| | # Items | 4,476 | 797 | 1,082 | 982 | 788 | 95 | 124 | 130 | 103 |
| | # Neuron-by-item pairs | 35,255 | 6,261 | 8,369 | 7,883 | 6,617 | 973 | 1,195 | 1,065 | 1,048 |
| | # Upper-tail neuron-by-item pairs | 83 | 22 | 37 | 16 | 8 | 5 | 3 | 5 | 4 |
| | # Upper-tail unique neurons | 65 | 22 | 31 | 14 | 7 | 5 | 3 | 5 | 4 |
| | **Amygdala** | | | | | | | | | |
| | # Items | 3,745 | 655 | 846 | 841 | 635 | 91 | 118 | 121 | 98 |
| | # Neuron-by-item pairs | 31,606 | 5,491 | 7,020 | 7,035 | 5,610 | 912 | 1,270 | 1,188 | 1,077 |
| | # Upper-tail neuron-by-item pairs | 74 | 21 | 14 | 13 | 14 | 1 | 2 | 0 | 5 |
| | # Upper-tail unique neurons | 53 | 19 | 12 | 11 | 13 | 1 | 2 | 0 | 5 |
| | **Anterior Cingulate Cortex** | | | | | | | | | |
| | # Items | 3,925 | 724 | 918 | 896 | 712 | 91 | 110 | 127 | 101 |
| | # Neuron-by-item pairs | 30,375 | 5,912 | 6,778 | 6,491 | 5,673 | 685 | 882 | 942 | 822 |
| | # Upper-tail neuron-by-item pairs | 73 | 14 | 16 | 11 | 13 | 1 | 3 | 2 | 5 |
| | # Upper-tail unique neurons | 59 | 13 | 15 | 11 | 13 | 1 | 3 | 2 | 5 |
| | **Prefrontal Cortex** | | | | | | | | | |
| | # Items | 3,318 | 593 | 815 | 721 | 576 | 68 | 90 | 98 | 84 |
| | # Neuron-by-item pairs | 22,873 | 4,033 | 5,098 | 4,702 | 4,023 | 672 | 787 | 804 | 859 |
| | # Upper-tail neuron-by-item pairs | 57 | 10 | 22 | 8 | 11 | 2 | 1 | 4 | 5 |
| | # Upper-tail unique neurons | 41 | 10 | 15 | 8 | 10 | 2 | 1 | 4 | 5 |

**Table S1.** Sampling density by retrieval demand and relative levels of spiking at encoding. Rows represent the number of trials, neuron-by-item pairs, upper-tail spike-count observations, and unique neurons contributing to the distributional analyses. Repeated items were categorized according to the relative levels of spiking activity immediately before and during encoding: relative increases in firing (Low-High), consistently elevated firing (High-High), relative decreases in firing (High-Low), and consistently low firing (Low-Low). Within each region, primary inferential analyses focused on remembered repeated item distributions from the High-High and Low-High encoding conditions, and the novel distribution. Novel trials combined correct rejections and false alarms, consistent with the repeated-versus-novel contrasts used throughout the manuscript. Neuron-by-item pairs correspond to the spike counts of one neuron in response to one item within each response distribution. Upper-tail neuron-by-item pairs reflect the number of responses exceeding the 0.9975 quantile of the pooled spike-count distribution within each brain region. Upper-tail unique neurons reflect the number of neurons contributing to at least one neuron-item pair above the 0.9975 quantile.

**Figure S1. Four moments of the remembered repeated and novel spike-count distributions across relative levels of spiking at encoding and brain regions**

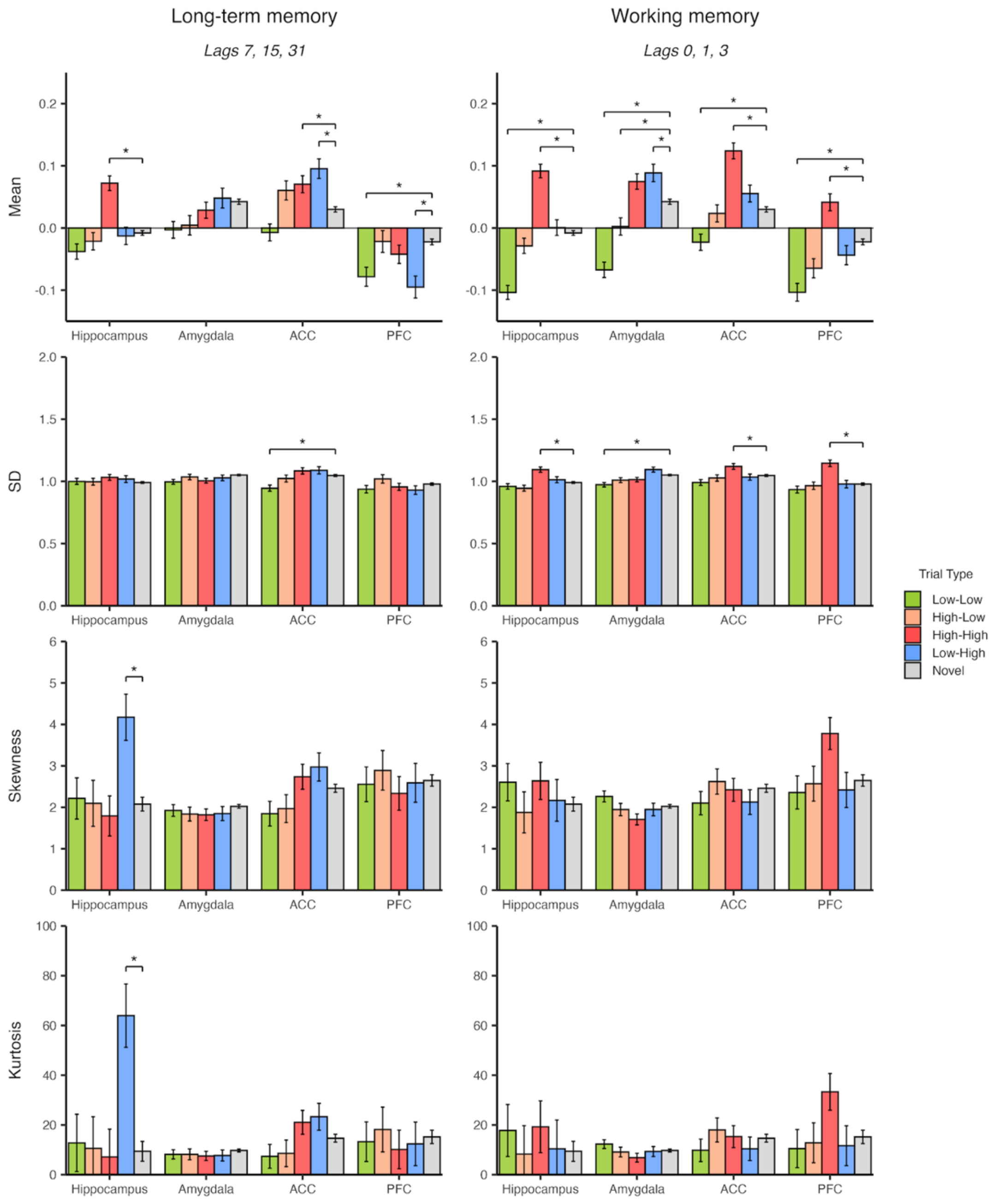


Mean, standard deviation, skewness, and kurtosis of normalized single-neuron spike-count distributions corresponding to recognition memory for remembered items. Each bar summarizes one moment of each normalized single-neuron spike-count distribution. The distributions consisted of neuron-by-item pairs representing the response of one neuron to one item. Four remembered item distributions each consisted of repeated items categorized according to the relative levels of spiking activity immediately before and during encoding: relative increases in firing (Low-High, blue), consistently elevated firing (High-High, red), relative decreases in firing (High-Low, orange), and consistently low firing (Low-Low, green). Graphs showing the mean and skewness for repeated items in the relative increase (Low-High) and consistently elevated (High-High) encoding conditions were reported in the main text but are included here for comparison. Error bars represent the standard deviation of the bootstrap distribution. Statistical comparisons were assessed using two-sided bootstrap tests (10,000 resamples). Brackets reflect comparisons that survived Benjamini-Hochberg false-discovery-rate correction. See Tables S2-S5 for full statistical reporting.

**Figure S2. Four moments of the forgotten repeated and novel spike-count distributions across relative levels of spiking at encoding and brain regions**

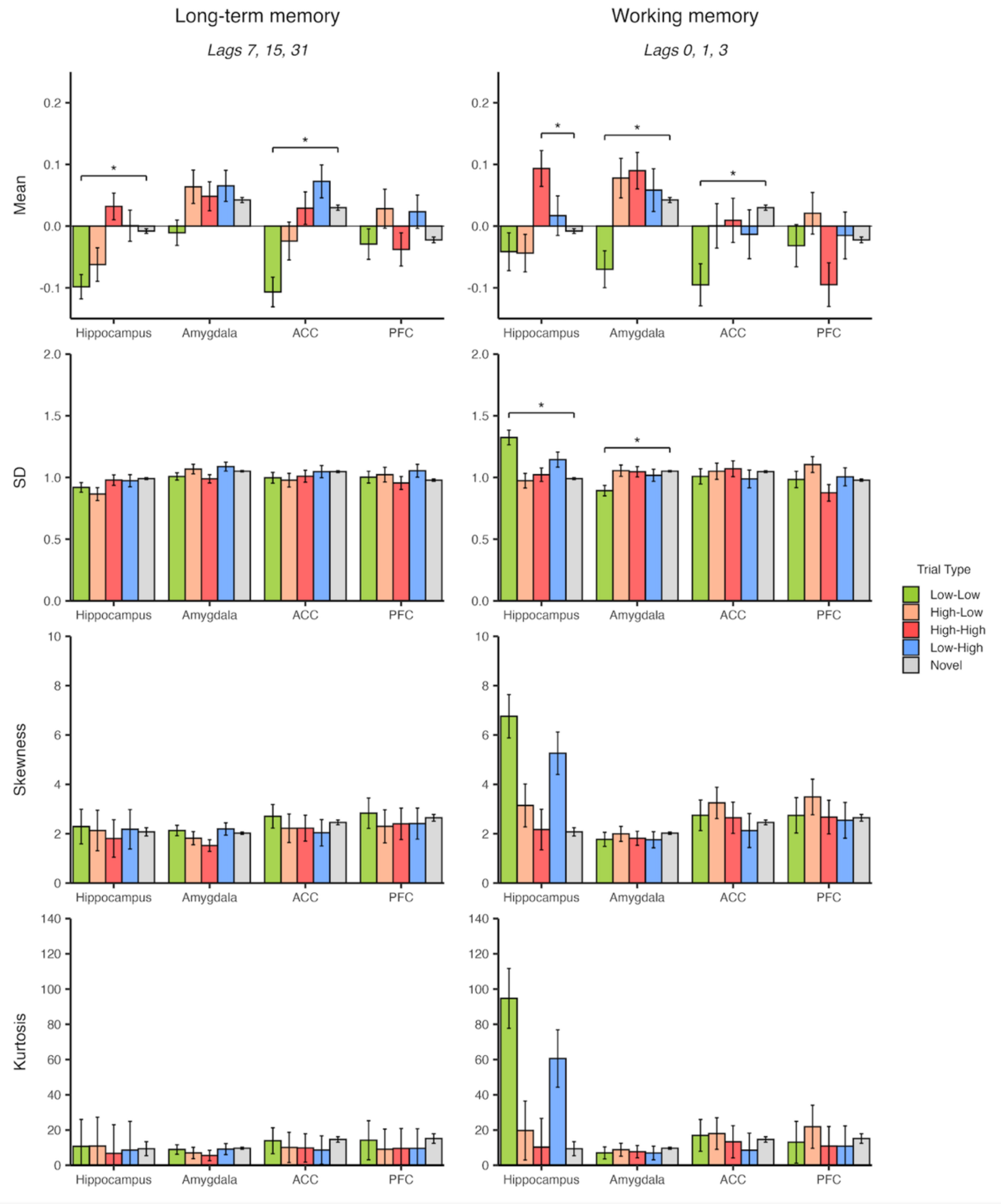


Mean, standard deviation, skewness, and kurtosis of normalized single-neuron spike-count distributions corresponding to recognition memory for forgotten items. Each bar summarizes one moment of each normalized single-neuron spike-count distribution. The distributions consisted of neuron-by-item pairs representing the response of one neuron to one item. Four forgotten item distributions each consisted of repeated items categorized according to the relative levels of spiking activity immediately before and during encoding: relative increases in firing (Low-High, blue), consistently elevated firing (High-High, red), relative decreases in firing (High-Low, orange), and consistently low firing (Low-Low, green). Error bars represent the standard deviation of the bootstrap distribution. Statistical comparisons were assessed using two-sided bootstrap tests (10,000 resamples). Brackets reflect comparisons that survived Benjamini-Hochberg false-discovery-rate correction. See Tables S2-S5 for full statistical reporting.

| Skewness | | | | | | | | | | | | | | | | | | | |
|---|---|---|---|---|---|---|---|---|---|---|---|---|---|---|---|---|---|---|---|
| **Remembered** | | | | | | | | | | | | | | | | | | | |
| | **Encoding** | **Hipp** | **Novel** | **p** | **ΔH-A** | **p(Int)** | **Amyg** | **Novel** | **p** | **ΔH-AC** | **p(Int)** | **ACC** | **Novel** | **p** | **ΔH-PF** | **p(Int)** | **PFC** | **Novel** | **p** |
| LTM | Low-High | 4.17 | 2.08 | **0.006** | 2.26 | **0.004** | 1.85 | 2.02 | 0.263 | 1.58 | **0.040** | 2.97 | 2.46 | 0.123 | 2.15 | **0.016** | 2.59 | 2.65 | 0.907 |
| | High-High | 1.79 | 2.08 | 0.505 | -0.09 | 0.854 | 1.82 | 2.02 | 0.138 | -0.57 | 0.284 | 2.74 | 2.46 | 0.377 | 0.02 | 0.955 | 2.34 | 2.65 | 0.428 |
| | High-Low | 2.09 | 2.08 | 0.972 | 0.19 | 0.659 | 1.84 | 2.02 | 0.231 | 0.50 | 0.341 | 1.97 | 2.46 | 0.134 | -0.23 | 0.651 | 2.89 | 2.65 | 0.577 |
| | Low-Low | 2.22 | 2.08 | 0.764 | 0.24 | 0.586 | 1.92 | 2.02 | 0.469 | 0.75 | 0.181 | 1.85 | 2.46 | 0.044 | 0.23 | 0.643 | 2.56 | 2.65 | 0.839 |
| WM | Low-High | 2.17 | 2.08 | 0.855 | 0.16 | 0.709 | 1.95 | 2.02 | 0.582 | 0.42 | 0.403 | 2.13 | 2.46 | 0.280 | 0.32 | 0.534 | 2.42 | 2.65 | 0.585 |
| | High-High | 2.64 | 2.08 | 0.180 | 0.87 | 0.085 | 1.71 | 2.02 | 0.033 | 0.60 | 0.254 | 2.42 | 2.46 | 0.902 | -0.57 | 0.302 | 3.78 | 2.65 | 0.014 |
| | High-Low | 1.88 | 2.08 | 0.664 | -0.12 | 0.783 | 1.94 | 2.02 | 0.573 | -0.36 | 0.474 | 2.62 | 2.46 | 0.622 | -0.12 | 0.790 | 2.57 | 2.65 | 0.861 |
| | Low-Low | 2.61 | 2.08 | 0.196 | 0.29 | 0.5 | 2.26 | 2.02 | 0.078 | 0.89 | 0.118 | 2.10 | 2.46 | 0.220 | 0.82 | 0.188 | 2.36 | 2.65 | 0.466 |
| **Forgotten** | | | | | | | | | | | | | | | | | | | |
| | **Encoding** | **Hipp** | **Novel** | **p** | **ΔH-A** | **p(Int)** | **Amyg** | **Novel** | **p** | **ΔH-AC** | **p(Int)** | **ACC** | **Novel** | **p** | **ΔH-PF** | **p(Int)** | **PFC** | **Novel** | **p** |
| LTM | Low-High | 2.18 | 2.08 | 0.874 | -0.07 | 0.898 | 2.19 | 2.02 | 0.44 | 0.52 | 0.382 | 2.04 | 2.46 | 0.404 | 0.34 | 0.529 | 2.41 | 2.65 | 0.662 |
| | High-High | 1.81 | 2.08 | 0.624 | 0.23 | 0.656 | 1.52 | 2.02 | 0.028 | -0.04 | 0.944 | 2.23 | 2.46 | 0.682 | -0.02 | 0.961 | 2.40 | 2.65 | 0.645 |
| | High-Low | 2.13 | 2.08 | 0.941 | 0.25 | 0.656 | 1.82 | 2.02 | 0.388 | 0.29 | 0.605 | 2.22 | 2.46 | 0.698 | 0.40 | 0.473 | 2.30 | 2.65 | 0.508 |
| | Low-Low | 2.29 | 2.08 | 0.698 | 0.10 | 0.833 | 2.13 | 2.02 | 0.578 | -0.04 | 0.947 | 2.71 | 2.46 | 0.630 | 0.03 | 0.951 | 2.83 | 2.65 | 0.729 |
| WM | Low-High | 5.26 | 2.08 | 0.015 | 3.44 | **0.013** | 1.76 | 2.02 | 0.334 | 3.51 | **0.018** | 2.13 | 2.46 | 0.629 | 3.29 | **0.037** | 2.54 | 2.65 | 0.884 |
| | High-High | 2.17 | 2.08 | 0.892 | 0.29 | 0.601 | 1.82 | 2.02 | 0.406 | -0.10 | 0.870 | 2.65 | 2.46 | 0.780 | 0.07 | 0.903 | 2.67 | 2.65 | 0.966 |
| | High-Low | 3.15 | 2.08 | 0.059 | 1.10 | 0.086 | 1.99 | 2.02 | 0.921 | 0.28 | 0.636 | 3.25 | 2.46 | 0.131 | 0.23 | 0.679 | 3.49 | 2.65 | 0.085 |
| | Low-Low | 6.76 | 2.08 | 0.014 | 4.93 | **0.010** | 1.77 | 2.02 | 0.307 | 4.39 | **0.010** | 2.75 | 2.46 | 0.647 | 4.59 | **0.012** | 2.74 | 2.65 | 0.879 |

**Table S2.** Item-specific retrieval signal (skewness) across relative levels of spiking at encoding and brain regions in the continuous recognition task for remembered and forgotten items. Repeated and novel distributions consisted of normalized single-neuron spike counts, with each observation representing the response of one neuron to one item. Repeated items were categorized according to the relative levels of spiking activity immediately before and during encoding: relative decreases in firing (High-Low) or consistently low firing (Low-Low). Values for items associated with a relative increase (Low-High) and consistently elevated (High-High) encoding firing were reported in Table 1 but are reported again here for comparison. Retrieval demand was defined by lag, with long-term memory (LTM) corresponding to lags 7, 15, and 31 and working memory (WM) corresponding to lags 0, 1, and 3. ΔH-Region denotes the difference between the hippocampal repeated-novel effect and the corresponding effect in the comparison region. Statistical comparisons were assessed using two-sided bootstrap tests (10,000 resamples). Bold *P* values survived Benjamini-Hochberg false-discovery-rate correction.

| Mean | | | | | | | | | | | | | | | | | | | |
|---|---|---|---|---|---|---|---|---|---|---|---|---|---|---|---|---|---|---|---|
| **Remembered** | | | | | | | | | | | | | | | | | | | |
| | **Encoding** | **Hipp** | **Novel** | **p** | **ΔH-A** | **p(Int)** | **Amyg** | **Novel** | **p** | **ΔH-AC** | **p(Int)** | **ACC** | **Novel** | **p** | **ΔH-PF** | **p(Int)** | **PFC** | **Novel** | **p** |
| LTM | Low-High | -0.01 | -0.01 | 0.741 | -0.01 | 0.628 | 0.05 | 0.04 | 0.728 | -0.07 | **0.001** | 0.10 | 0.03 | **<0.001** | 0.07 | **0.004** | -0.10 | -0.02 | **<0.001** |
| | High-High | 0.07 | -0.01 | **<0.001** | 0.09 | **<0.001** | 0.03 | 0.04 | 0.312 | 0.04 | **0.037** | 0.07 | 0.03 | **0.006** | 0.10 | **<0.001** | -0.04 | -0.02 | 0.199 |
| | High-Low | -0.02 | -0.01 | 0.349 | 0.02 | 0.268 | 0.00 | 0.04 | 0.019 | -0.04 | 0.040 | 0.06 | 0.03 | 0.055 | -0.01 | 0.543 | -0.02 | -0.02 | 0.976 |
| | Low-Low | -0.04 | -0.01 | 0.022 | 0.02 | 0.422 | 0.00 | 0.04 | 0.002 | 0.01 | 0.702 | -0.01 | 0.03 | 0.010 | 0.03 | 0.201 | -0.08 | -0.02 | **<0.001** |
| WM | Low-High | 0.001 | -0.01 | 0.515 | -0.04 | 0.057 | 0.09 | 0.04 | **0.001** | -0.02 | 0.376 | 0.06 | 0.03 | 0.073 | 0.03 | 0.148 | -0.04 | -0.02 | 0.182 |
| | High-High | 0.09 | -0.01 | **<0.001** | 0.07 | **<0.001** | 0.07 | 0.04 | 0.014 | 0.01 | 0.762 | 0.12 | 0.03 | **<0.001** | 0.04 | 0.053 | 0.04 | -0.02 | **<0.001** |
| | High-Low | -0.03 | -0.01 | 0.105 | 0.02 | 0.326 | 0.00 | 0.04 | **0.005** | -0.01 | 0.453 | 0.02 | 0.03 | 0.653 | 0.02 | 0.293 | -0.06 | -0.02 | 0.009 |
| | Low-Low | -0.10 | -0.01 | **<0.001** | 0.01 | 0.432 | -0.07 | 0.04 | **<0.001** | -0.04 | **0.016** | -0.02 | 0.03 | **<0.001** | -0.01 | 0.452 | -0.10 | -0.02 | **<0.001** |
| **Forgotten** | | | | | | | | | | | | | | | | | | | |
| | **Encoding** | **Hipp** | **Novel** | **p** | **ΔH-A** | **p(Int)** | **Amyg** | **Novel** | **p** | **ΔH-AC** | **p(Int)** | **ACC** | **Novel** | **p** | **ΔH-PF** | **p(Int)** | **PFC** | **Novel** | **p** |
| LTM | Low-High | 0.001 | -0.01 | 0.730 | -0.01 | 0.694 | 0.07 | 0.04 | 0.375 | -0.03 | 0.373 | 0.07 | 0.03 | 0.113 | -0.04 | 0.327 | 0.02 | -0.02 | 0.097 |
| | High-High | 0.03 | -0.01 | 0.070 | 0.03 | 0.297 | 0.05 | 0.04 | 0.796 | 0.04 | 0.239 | 0.03 | 0.03 | 0.970 | 0.06 | 0.110 | -0.04 | -0.02 | 0.567 |
| | High-Low | -0.06 | -0.01 | 0.045 | -0.08 | 0.051 | 0.06 | 0.04 | 0.431 | 0.00 | 0.995 | -0.02 | 0.03 | 0.078 | -0.10 | **0.012** | 0.03 | -0.02 | 0.111 |
| | Low-Low | -0.10 | -0.01 | **<0.001** | -0.04 | 0.196 | -0.01 | 0.04 | 0.011 | 0.05 | 0.143 | -0.11 | 0.03 | **<0.001** | -0.08 | **0.009** | -0.03 | -0.02 | 0.789 |
| WM | Low-High | 0.02 | -0.01 | 0.443 | 0.01 | 0.848 | 0.06 | 0.04 | 0.643 | 0.07 | 0.191 | -0.01 | 0.03 | 0.278 | 0.02 | 0.720 | -0.02 | -0.02 | 0.852 |
| | High-High | 0.09 | -0.01 | **<0.001** | 0.05 | 0.188 | 0.09 | 0.04 | 0.107 | 0.12 | **0.009** | 0.01 | 0.03 | 0.564 | 0.17 | **<0.001** | -0.09 | -0.02 | 0.039 |
| | High-Low | -0.04 | -0.01 | 0.245 | -0.07 | 0.109 | 0.08 | 0.04 | 0.277 | -0.01 | 0.898 | 0.00 | 0.03 | 0.419 | -0.08 | 0.086 | 0.02 | -0.02 | 0.206 |
| | Low-Low | -0.04 | -0.01 | 0.280 | 0.08 | 0.069 | -0.07 | 0.04 | **<0.001** | 0.09 | 0.046 | -0.10 | 0.03 | **<0.001** | -0.02 | 0.606 | -0.03 | -0.02 | 0.788 |

**Table S3.** Generic recognition signal (mean) across relative levels of spiking at encoding and brain regions in the continuous recognition task for remembered and forgotten items. Repeated and novel distributions consisted of normalized single-neuron spike counts, with each observation representing the response of one neuron to one item. Repeated items were categorized according to the relative levels of spiking activity immediately before and during encoding: relative decreases in firing (High-Low) or consistently low firing (Low-Low). Values for items associated with a relative increase (Low-High) and consistently elevated (High-High) encoding firing were reported in Table 2 but are reported again here for comparison. Retrieval demand was defined by lag, with long-term memory (LTM) corresponding to lags 7, 15, and 31 and working memory (WM) corresponding to lags 0, 1, and 3. ΔH-Region denotes the difference between the hippocampal repeated-novel effect and the corresponding effect in the comparison region. Statistical comparisons were assessed using two-sided bootstrap tests (10,000 resamples). Bold *P* values survived Benjamini-Hochberg false-discovery-rate correction.

| Kurtosis | | | | | | | | | | | | | | | | | | | |
|---|---|---|---|---|---|---|---|---|---|---|---|---|---|---|---|---|---|---|---|
| **Remembered** | | | | | | | | | | | | | | | | | | | |
| | **Encoding** | **Hipp** | **Novel** | **p** | **ΔH-A** | **p(Int)** | **Amyg** | **Novel** | **p** | **ΔH-AC** | **p(Int)** | **ACC** | **Novel** | **p** | **ΔH-PF** | **p(Int)** | **PFC** | **Novel** | **p** |
| LTM | Low-High | 63.94 | 9.40 | **0.003** | 56.50 | **0.003** | 7.74 | 9.71 | 0.239 | 45.90 | **0.014** | 23.30 | 14.66 | 0.092 | 57.35 | **0.004** | 12.38 | 15.20 | 0.734 |
| | High-High | 7.14 | 9.40 | 0.814 | -0.09 | 0.993 | 7.54 | 9.71 | 0.151 | -8.64 | 0.367 | 21.04 | 14.66 | 0.179 | 2.78 | 0.749 | 10.15 | 15.20 | 0.475 |
| | High-Low | 10.60 | 9.40 | 0.907 | 2.71 | 0.769 | 8.19 | 9.71 | 0.376 | 7.28 | 0.439 | 8.58 | 14.66 | 0.227 | -1.76 | 0.831 | 18.15 | 15.20 | 0.727 |
| | Low-Low | 12.78 | 9.40 | 0.713 | 4.90 | 0.576 | 8.18 | 9.71 | 0.316 | 10.66 | 0.289 | 7.37 | 14.66 | 0.121 | 5.31 | 0.557 | 13.26 | 15.20 | 0.817 |
| WM | Low-High | 10.40 | 9.40 | 0.924 | 1.40 | 0.879 | 9.31 | 9.71 | 0.803 | 5.24 | 0.569 | 10.42 | 14.66 | 0.386 | 4.55 | 0.614 | 11.64 | 15.20 | 0.641 |
| | High-High | 19.26 | 9.40 | 0.249 | 12.72 | 0.173 | 6.84 | 9.71 | 0.077 | 9.21 | 0.347 | 15.31 | 14.66 | 0.906 | -8.24 | 0.408 | 33.30 | 15.20 | 0.037 |
| | High-Low | 8.29 | 9.40 | 0.913 | -0.51 | 0.955 | 9.10 | 9.71 | 0.712 | -4.44 | 0.624 | 17.99 | 14.66 | 0.523 | 1.26 | 0.878 | 12.82 | 15.20 | 0.762 |
| | Low-Low | 17.78 | 9.40 | 0.321 | 5.79 | 0.512 | 12.30 | 9.71 | 0.093 | 13.23 | 0.2 | 9.80 | 14.66 | 0.294 | 13.09 | 0.257 | 10.48 | 15.20 | 0.511 |
| **Forgotten** | | | | | | | | | | | | | | | | | | | |
| | **Encoding** | **Hipp** | **Novel** | **p** | **ΔH-A** | **p(Int)** | **Amyg** | **Novel** | **p** | **ΔH-AC** | **p(Int)** | **ACC** | **Novel** | **p** | **ΔH-PF** | **p(Int)** | **PFC** | **Novel** | **p** |
| LTM | Low-High | 8.62 | 9.40 | 0.959 | -0.24 | 0.982 | 9.16 | 9.71 | 0.842 | 5.23 | 0.529 | 8.65 | 14.66 | 0.381 | 4.77 | 0.545 | 9.64 | 15.20 | 0.486 |
| | High-High | 6.79 | 9.40 | 0.825 | 1.53 | 0.881 | 5.56 | 9.71 | 0.058 | 2.19 | 0.772 | 9.86 | 14.66 | 0.527 | 2.98 | 0.696 | 9.59 | 15.20 | 0.482 |
| | High-Low | 10.94 | 9.40 | 0.920 | 4.22 | 0.687 | 7.03 | 9.71 | 0.233 | 6.04 | 0.465 | 10.16 | 14.66 | 0.601 | 7.65 | 0.360 | 9.08 | 15.20 | 0.443 |
| | Low-Low | 10.78 | 9.40 | 0.917 | 2.13 | 0.829 | 8.96 | 9.71 | 0.736 | 2.10 | 0.791 | 13.94 | 14.66 | 0.942 | 2.36 | 0.762 | 14.22 | 15.20 | 0.925 |
| WM | Low-High | 60.60 | 9.40 | 0.018 | 53.92 | **0.015** | 6.99 | 9.71 | 0.286 | 57.30 | **0.021** | 8.56 | 14.66 | 0.466 | 55.52 | **0.039** | 10.87 | 15.20 | 0.677 |
| | High-High | 10.37 | 9.40 | 0.949 | 2.95 | 0.778 | 7.72 | 9.71 | 0.430 | 2.29 | 0.759 | 13.33 | 14.66 | 0.908 | 5.23 | 0.496 | 10.93 | 15.20 | 0.667 |
| | High-Low | 19.74 | 9.40 | 0.264 | 11.21 | 0.227 | 8.83 | 9.71 | 0.769 | 6.98 | 0.402 | 18.01 | 14.66 | 0.767 | 3.66 | 0.633 | 21.87 | 15.20 | 0.406 |
| | Low-Low | 94.71 | 9.40 | 0.016 | 87.99 | **0.014** | 7.03 | 9.71 | 0.255 | 83.02 | **0.011** | 16.96 | 14.66 | 0.834 | 87.43 | **0.014** | 13.08 | 15.20 | 0.859 |

**Table S4.** Kurtosis of normalized single-neuron spike-count distributions across relative levels of spiking at encoding and brain regions. Repeated and novel distributions consisted of normalized single-neuron spike counts, with each observation representing the response of one neuron to one item. Repeated items were categorized according to the relative levels of spiking activity immediately before and during encoding: relative decreases in firing (High-Low) or consistently low firing (Low-Low). Retrieval demand was defined by lag, with long-term memory (LTM) corresponding to lags 7, 15, and 31 and working memory (WM) corresponding to lags 0, 1, and 3. ΔH-Region denotes the difference between the hippocampal repeated-novel effect and the corresponding effect in the comparison region. Statistical comparisons were assessed using two-sided bootstrap tests (10,000 resamples). Bold *P* values survived Benjamini-Hochberg false-discovery-rate correction.

| Standard Deviation | | | | | | | | | | | | | | | | | | | |
|---|---|---|---|---|---|---|---|---|---|---|---|---|---|---|---|---|---|---|---|
| **Remembered** | | | | | | | | | | | | | | | | | | | |
| | **Encoding** | **Hipp** | **Novel** | **p** | **ΔH-A** | **p(Int)** | **Amyg** | **Novel** | **p** | **ΔH-AC** | **p(Int)** | **ACC** | **Novel** | **p** | **ΔH-PF** | **p(Int)** | **PFC** | **Novel** | **p** |
| LTM | Low-High | 1.02 | 0.99 | 0.306 | 0.05 | 0.159 | 1.03 | 1.05 | 0.336 | -0.01 | 0.737 | 1.09 | 1.05 | 0.163 | 0.08 | 0.099 | 0.93 | 0.98 | 0.169 |
| | High-High | 1.03 | 0.99 | 0.09 | 0.09 | **0.006** | 1.00 | 1.05 | 0.02 | 0.00 | 0.906 | 1.08 | 1.05 | 0.157 | 0.06 | 0.104 | 0.96 | 0.98 | 0.453 |
| | High-Low | 1.00 | 0.99 | 0.817 | 0.02 | 0.547 | 1.04 | 1.05 | 0.5 | 0.03 | 0.435 | 1.02 | 1.05 | 0.397 | -0.04 | 0.437 | 1.02 | 0.98 | 0.238 |
| | Low-Low | 1.00 | 0.99 | 0.699 | 0.06 | 0.049 | 1.00 | 1.05 | 0.006 | 0.11 | **0.003** | 0.95 | 1.05 | **<0.001** | 0.05 | 0.208 | 0.94 | 0.98 | 0.189 |
| WM | Low-High | 1.01 | 0.99 | 0.369 | -0.02 | 0.515 | 1.09 | 1.05 | 0.038 | 0.04 | 0.33 | 1.03 | 1.05 | 0.623 | 0.02 | 0.578 | 0.98 | 0.98 | 0.997 |
| | High-High | 1.09 | 0.99 | **<0.001** | 0.14 | **<0.001** | 1.01 | 1.05 | 0.046 | 0.03 | 0.358 | 1.12 | 1.05 | **0.004** | -0.06 | 0.078 | 1.15 | 0.98 | **<0.001** |
| | High-Low | 0.95 | 0.99 | 0.068 | 0.00 | 0.899 | 1.01 | 1.05 | 0.046 | -0.03 | 0.48 | 1.03 | 1.05 | 0.451 | -0.03 | 0.412 | 0.96 | 0.98 | 0.666 |
| | Low-Low | 0.96 | 0.99 | 0.166 | 0.05 | 0.113 | 0.97 | 1.05 | **<0.001** | 0.03 | 0.458 | 0.99 | 1.05 | 0.025 | 0.01 | 0.731 | 0.93 | 0.98 | 0.127 |
| **Forgotten** | | | | | | | | | | | | | | | | | | | |
| | **Encoding** | **Hipp** | **Novel** | **p** | **ΔH-A** | **p(Int)** | **Amyg** | **Novel** | **p** | **ΔH-AC** | **p(Int)** | **ACC** | **Novel** | **p** | **ΔH-PF** | **p(Int)** | **PFC** | **Novel** | **p** |
| LTM | Low-High | 0.97 | 0.99 | 0.700 | -0.05 | 0.354 | 1.09 | 1.05 | 0.309 | -0.02 | 0.800 | 1.05 | 1.05 | 0.995 | -0.09 | 0.182 | 1.05 | 0.98 | 0.132 |
| | High-High | 0.98 | 0.99 | 0.766 | 0.05 | 0.326 | 0.99 | 1.05 | 0.070 | 0.03 | 0.677 | 1.01 | 1.05 | 0.439 | 0.01 | 0.864 | 0.96 | 0.98 | 0.659 |
| | High-Low | 0.87 | 0.99 | 0.027 | -0.14 | 0.036 | 1.07 | 1.05 | 0.669 | -0.06 | 0.438 | 0.98 | 1.05 | 0.207 | -0.17 | 0.043 | 1.02 | 0.98 | 0.431 |
| | Low-Low | 0.92 | 0.99 | 0.069 | -0.03 | 0.541 | 1.01 | 1.05 | 0.149 | -0.02 | 0.714 | 1.00 | 1.05 | 0.256 | -0.10 | 0.124 | 1.00 | 0.98 | 0.600 |
| WM | Low-High | 1.14 | 0.99 | 0.023 | 0.19 | **0.022** | 1.02 | 1.05 | 0.494 | 0.21 | **0.031** | 0.99 | 1.05 | 0.397 | 0.13 | 0.170 | 1.01 | 0.98 | 0.689 |
| | High-High | 1.02 | 0.99 | 0.521 | 0.04 | 0.581 | 1.05 | 1.05 | 0.908 | 0.01 | 0.915 | 1.07 | 1.05 | 0.696 | 0.13 | 0.119 | 0.88 | 0.98 | 0.115 |
| | High-Low | 0.97 | 0.99 | 0.747 | -0.02 | 0.762 | 1.06 | 1.05 | 0.933 | -0.02 | 0.808 | 1.05 | 1.05 | 0.960 | -0.14 | 0.094 | 1.10 | 0.98 | 0.054 |
| | Low-Low | 1.32 | 0.99 | **<0.001** | 0.49 | **<0.001** | 0.89 | 1.05 | **<0.001** | 0.37 | **<0.001** | 1.01 | 1.05 | 0.524 | 0.33 | **0.001** | 0.98 | 0.98 | 0.924 |

**Table S5.** Standard deviation of normalized single-neuron spike-count distributions across relative levels of spiking at encoding and brain regions. Repeated and novel distributions consisted of normalized single-neuron spike counts, with each observation representing the response of one neuron to one item. Repeated items were categorized according to the relative levels of spiking activity immediately before and during encoding: relative increases in firing (Low-High), consistently elevated firing (High-High), relative decreases in firing (High-Low), or consistently low firing (Low-Low). Retrieval demand was defined by lag, with long-term memory (LTM) corresponding to lags 7, 15, and 31 and working memory (WM) corresponding to lags 0, 1, and 3. ΔH-Region denotes the difference between the hippocampal repeated-novel effect and the corresponding effect in the comparison region. Statistical comparisons were assessed using two-sided bootstrap tests (10,000 resamples). Bold *P* values survived Benjamini-Hochberg false-discovery-rate correction.
